# scDRP: Disentangled representation learning for predicting single-cell responses to perturbations and estimating individual treatment effects

**DOI:** 10.1101/2025.11.21.689783

**Authors:** Jianle Sun, Petar Stojanov, Kun Zhang

**Affiliations:** Carnegie Mellon University, Pittsburgh, 15213, PA, United States of America; Broad institute of MIT and Harvard, Cambridge, 02142, MA, United States of America; Mohamed bin Zayed University of Artificial Intelligence, Masdar City Abu Dhabi, United Arab Emirates

**Keywords:** single-cell perturbation, individual treatment effect, disentangled representation, optimal transport, heterogeneous cellular response, functional gene module

## Abstract

Dissecting cell-state-specific changes in gene regulation induced by perturbations is crucial for understanding biological mechanisms. However, single-cell sequencing provides only unmatched snapshots of cells under different conditions. This destructive measurement process hinders the estimation of individualized treatment effects (ITEs), which are essential for pinpointing these heterogeneous mechanistic responses. We develop scDRP, a generative framework that leverages disentangled representation learning with asymptotic correctness guarantees to separate perturbation-induced and perturbation-invariant latent variables via a sparsity regularized ***β***-VAE. Assuming quantile-preserving effects of perturbations conditional on confounders, scDRP performs conditional optimal transport in the disentangled latent space to infer counterfactual states and estimate ITEs. Applied to simulated and real single-cell perturbation data, scDRP accurately estimates treatment effects and individual counterfactual responses, with subsequent biclustering analysis further elucidating cell-type-specific functional gene module dynamics. Specifically, it captures distinct cellular patterns under rhinovirus and cigarette-smoke extract exposures, reveals heterogeneous responses to interferon stimulation across diverse immune cell types, and identifies distinct functional module activation in chronic myeloid leukemia cells following CRISPR activation targeting different genes. scDRP also generalizes to unseen perturbation doses and combinations. Our framework provides a principled computational approach to extracting heterogeneous causal relationships from single-cell perturbation data, enabling a deeper understanding of cellular and molecular mechanisms.

## 1 Introduction

Understanding cell-specific heterogeneous responses to chemical or genetic perturbations is crucial to deciphering the principles of life. Advances in single-cell sequencing technologies, such as Perturb-seq [1] and sci-Plex [2], now enable large-scale profiling of multi-omics features across numerous perturbations, offering an unprecedented opportunity to dissect gene regulatory circuitry in development and disease [3]. However, the measurements consist of unmatched populations of cells under different perturbations, and this destructive nature of sequencing experiments precludes direct comparison of a cell’s state before and after perturbation, which is formally referred to as the individual treatment effect (ITE), posing a fundamental challenge for causal inference.

To circumvent this, many prevailing perturbation modeling frameworks, such as scGEN [4] and State [5], focus on predicting population-average responses; although effective at the population level, they obscure treatment heterogeneity, which is crucial in complex systems like organoids, where subpopulations respond differently. Furthermore, typical methods aiming for individual-level prediction often resort to a branch of strong assumptions. CINEMA-OT [6], for instance, assumes that perturbation-induced latent signals are completely independent of confounders (the cell’s intrinsic properties such as cell types), and that responses arise from linear combinations, neglecting the complex nonlinear relationships and interactions that govern cellular responses.

In this paper, to address this challenge, we propose a framework of single-cell disentangled representation learning with perturbations (scDRP). Leveraging the properties of genetic perturbations and recent advances in disentangled representation learning, scDRP employs a well-designed sparsity-regularized *β*-VAE to learn and separate perturbation-dependent and perturbation-independent latent variables. Under the assumption that perturbations exert quantile-preserving effects on the latent variables they influence when conditioning on confounders, our framework enables the identification of ITE. Biologically, this assumption implies that a cell’s intrinsic relative rank within a homogeneous subpopulation (e.g., within the same cell type, with similar cell states) is preserved after treatments. For instance, a cell that is inherently the most sensitive to the perturbation (e.g., 95th percentile) among its control peers is assumed to remain the most sensitive (95th percentile) relative to the new perturbed population, even as the entire distribution shifts. Specifically, conditional optimal transport (OT) with squared Euclidean cost, which is conceptually equivalent to quantile matching or monotone rearrangement in one-dimension [7], is performed in the disentangled latent space to generate counterfactual predictions and estimate individual causal responses (Fig. 1a).

**Fig. 1.**
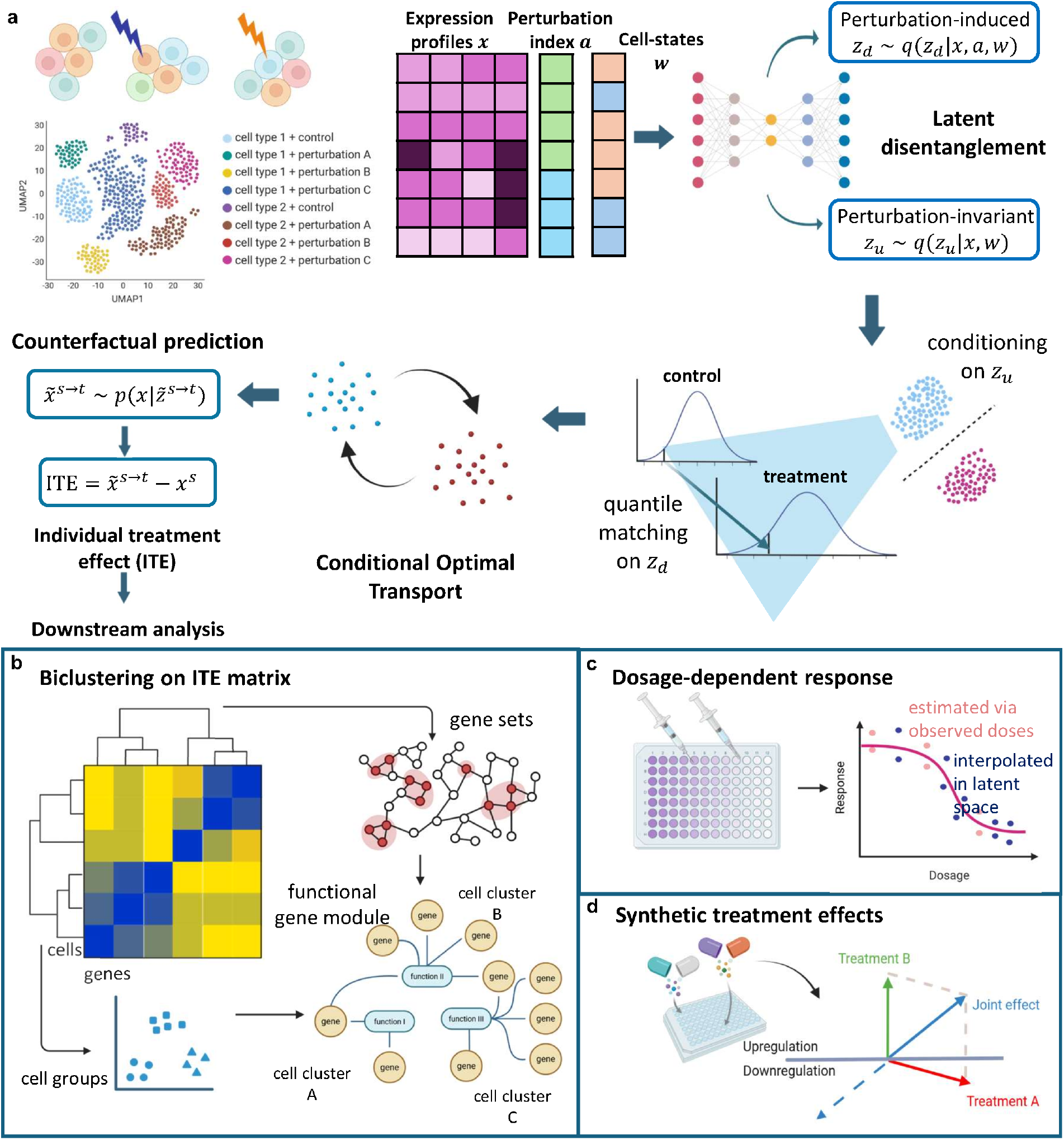
Model overview. **a**. The fundamental idea of scDRP is that the counterfactual twin of a control cell should be a cell within the perturbed group that shares a similar intrinsic cell identity and maintains its relative position (rank) along the perturbation response axis. scDRP first disentangles the latent space into perturbation-dependent cell responses (***z***_*d*_) and perturbation-independent cell identities (***z***_*u*_). By performing quantile matching on ***z***_*d*_ conditional on ***z***_*u*_ (implemented via convex-cost OT with a soft penalty on conditionals), scDRP identifies individualized causal effects and formulates them as a conditional optimal transport problem in the latent space. **b**. Biclustering of the estimated ITE matrix reveals functional gene modules underlying perturbation responses across different conditions and cell populations. **c**. scDRP estimates effects of unseen dosages by interpolation in the latent space. **d**. scDRP infers unseen combinatorial perturbation effects by vector addition of the estimated ITEs.

We demonstrate the strong performance of scDRP in estimating ITEs and predicting counterfactual perturbation responses across both simulated and real single-cell perturbation datasets. Biclustering analysis on the estimated ITE matrix reveals cell type-specific differences in functional gene module (FGM) responses under diverse perturbations (Fig. 1b). Specifically, the re-analysis of rhinovirus and cigarette-smoke exposure data resolves distinct, fine-grained cellular patterns. Applying to additional datasets, we further uncover heterogeneous responses to interferon stimulation across diverse immune cell types, and delineate distinct functional module activation in chronic myeloid leukemia cells with targeted CRISPR activation (CRISPRa) Perturb-seq. Furthermore, by performing interpolation within the latent and effect space, we enable inference of the effects of unobserved perturbation doses (Fig. 1c) as well as combinations of multiple perturbations (Fig. 1d), highlighting the model’s ability to generalize to unseen scenarios.

Our theoretical analysis and empirical results suggest that scDRP will provide a powerful computational tool for dissecting large-scale single-cell perturbation datasets, advancing our understanding of cellular and molecular mechanisms underlying physiological and disease processes and paving the way for systematic *in silico* perturbation screening and rational therapeutic target discovery.

## 2 Results

### 2.1 Overview of the scDRP framework for inferring single-cell treatment effects

To overcome the fundamental limitation that single-cell sequencing cannot capture the same cells across control and perturbed states, we developed scDRP, a deep generative framework designed to infer individual treatment effects from unmatched population data. Implemented as Python package (https://github.com/sjl-sjtu/scDRP), scDRP takes as input a single-cell expression matrix (log-normalized values or raw counts), perturbation index for each cell, and, optionally, covariates such as cell-type labels or sequencing batch. From these it estimates how each control cell would have responded to the given perturbations, i.e., the ITE matrix, and further aggregates these into biologically interpretable functional gene modules.

Intuitively, the transcriptional state of a cell can be viewed as comprising two parts: who the cell is (its stable, intrinsic identity, such as cell type and maturation stage) and how it responds to external stimuli. Because of the disruptive nature of single-cell sequencing, we can never observe the same cell before and after a perturbation, so we cannot directly read off a cell’s response. scDRP addresses this by separating the two parts in a learned latent space, holding identity fixed, and asking what the response part would look like under the perturbation. The matching between an untreated cell and its perturbed counterpart relies on a natural principle: a perturbation may shift the whole population, but a cell’s relative position (i.e., rank or percentile) within cells like itself is preserved. Aligning these ranks, which OT performs automatically, yields an *in silico* “counterfactual twin” for each cell, and hence its individual treatment effect.

To implement this conceptual framework (Fig. 1a), scDRP posits a generative model where the high-dimensional gene expression profile ***x*** is a noisy function of low-dimensional latent representation ***z***. We structured this latent space into two distinct conditionally independent subspaces: perturbation-invariant components ***z***_*u*_ and perturbation-induced components ***z***_*d*_. Biologically, ***z***_*u*_ captures intrinsic cellular properties (convoluted with confounders ***w***) that remain stable across conditions. In contrast, ***z***_*d*_ captures the variation induced by cellular responses to perturbations, which is jointly affected by the perturbation ***a*** and confounders ***w***.

Technically, we operationalize this using a *β*-Variational Autoencoder (*β*-VAE) regularized with a Hilbert–Schmidt Conditional Independence Criterion (HSCIC) [8, 9] and gated L_0_ norm [10]. This regularization enforces strict conditional independence between ***z***_*u*_ and ***z***_*d*_ given ***w***, ensuring that the decomposed “identity” features do not leak into the “response” space (see Supplementary Notes SN1 for identifiability discussion [11, 12]).

With the disentangled latent space, scDRP estimates ITEs by mapping cells from the control to the treated distribution. A key challenge in causal inference is determining which control cell corresponds to which treated cell. We address this by assuming that perturbations exert rank-preserving effects on the response latent variables ***z***_*d*_ within homogeneous subpopulations (i.e., after controlling for confounders ***w*** or its surrogate ***z***_*u*_). This corresponds to the conservative coupling that minimizes the overall ITEs (i.e., minimal change principle) [13], implying that a cell’s relative sensitivity rank or quantile within its peer group is preserved after treatment.

To effectively conduct quantile matching within similar cells across different conditions, scDRP formulates it as a conditional OT problem as OT with a convex cost function is mathematically equivalent to quantile matching in 1-dimension [7]. We seek an optimal coupling matrix Γ^∗^ that matches control and treated cells by minimizing the transport cost in the response space (***z***_*d*_) while strictly constraining them to share similar intrinsic identities (***z***_*u*_) by imposing a soft conditional constraint. Mathematically, this is achieved by solving for the coupling that minimizes ∫*C*(***z***^*s*^, ***z***^*t*^)*d*Γ(***z***^*s*^, ***z***^*t*^), where a conditional-penalized cost

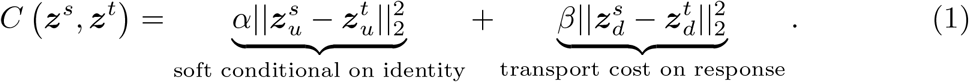

Then scDRP can generate counterfactual predictions and estimate ITE for each cell through barycentric projection with optimal coupling Γ^∗^ (Method & Supplementary Notes SN2). We also illustrate the idea of conditional OT in disentangled latent space with a toy example (in Supplementary Notes SN3 & Supplementary Fig. S1-S3). This formulation allows scDRP to move beyond population averages and resolve heterogeneity at single-cell resolution.

### 2.2 scDRP accurately estimates cell-specific individual treatment effects on different genes

Since the true counterfactual outcome of a cell is never observable in real destructive single-cell experiments, simulation is the only setting in which estimated ITEs and CATEs, as well as the recovered latent disentanglement and control–treatment matching, can be evaluated against ground truth; we therefore begin with simulated benchmarks before turning to real data. We compared our approach with two representative existing methods, CINEMA-OT (COT) [6] and scGEN [4]. CINEMA-OT assumes a linear mixing process and strictly independent perturbation variables (*z*_*d*_) from confounders. In contrast, scGEN models a nonlinear mapping but lacks latent space disentanglement, relying on average changes for inference. Our model advances both by incorporating a general nonlinear function with proper latent disentanglement. Furthermore, we relax the independence constraint, requiring *z*_*d*_ and *z*_*u*_ to be independent only conditional on confounders (e.g., cell type), which better aligns with realistic biological settings.

We generated simulated data comprising five cell types subjected to five perturbation conditions under varying dimensionalities of *z*_*d*_ and *z*_*u*_. Each setting was repeated 20 times to eliminate random variation. We estimated both the ITE and CATE under two settings: with known cell-type labels (observed confounding surrogates) and with unknown cell-type labels (unobserved confounding surrogates). We evaluated the average root mean square error (RMSE) of ITE and conditional average treatment effect (CATE) across all perturbations and all cell types. As shown in Fig. 2a, CINEMA-OT performs poorly on complex nonlinear data, partly because it relies on linear disentanglement, and partly due to its overly strong assumption that confounders are completely independent of the treatment. Although scGEN employs nonlinear neural networks and performs relatively well in CATE estimation through mean-shift–based adaptation instead of latent disentanglement, it performs poorly in ITE estimation. In contrast, our method achieves consistently accurate and stable ITE estimation in both settings.

**Fig. 2.**
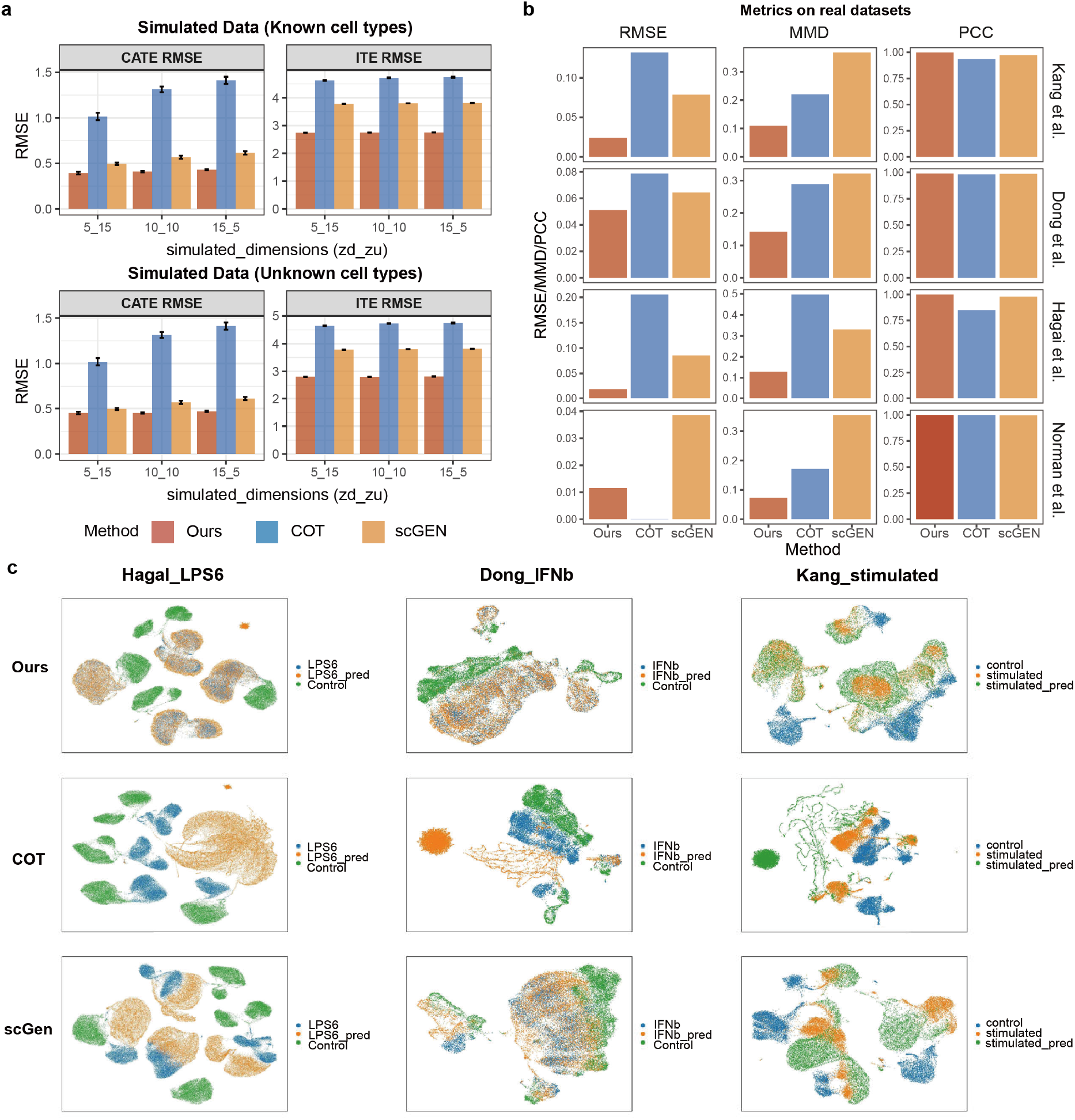
Performance of effect estimation and counterfactual prediction on simulated and real datasets. **a**. RMSE of CATE and ITE estimation on all genes under known and unknown cell type scenarios. The x-axis shows the dimension settings for *z*_*d*_ and *z*_*u*_ in the simulation, and the different colors correspond to different methods. We repeat each simulated setting 20 times and report the mean*±*1.96 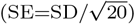. **b**. Performance on counterfactual predictions on real single-cell data, where evaluation metrics includes RMSE and PCC of the mean expression of each gene within top 3000 highly variable genes (HVGs) between observed treatment group and counterfactual predictions from control group (we compare the mean since the true counterfactual outcome of control cells are not accessible in real data), as well as MMD between the observed and predicted distributions. Each row shows results on one dataset and each column represents one metrics. All metrics are averaged across all perturbations and cell types. **c**. UMAP visualization of observed and predicted cells under counterfactual perturbation context, with blue scatters representing control-group cells, orange representing treatment-group cells, and green representing predicted responses of control-group cells under the treatment.

We also discuss different configurations for applying optimal transport and barycentric projection in the disentangled latent space. As shown in Supplementary Fig. S8, when cell types are known, the model is largely insensitive to hyperparameter choices and projection schemes, as the grouped optimal transport enables effective domain adaptation in the latent space. In contrast, when cell types are unknown, carefully balancing the weights between the conditional and transport cost components becomes more critical. Besides, the simulation can provide the ground-truth one-to-one pairing by simulating the same cell in both states that destructive sequencing cannot, allowing us to test the matching directly: the conditional OT reproduces this pairing exactly within every cell type on the ground-truth factors and largely preserves it on the model-inferred factors (Supplementary Note SN4, Fig. S4, Table S1).

### 2.3 scDRP effectively predicts cellular responses to diverse perturbations *in silico* on real datasets

We then evaluate the predictive capability on perturbation response by comparing the observed real treated samples with the pseudo-treated samples generated through *in silico* counterfactual prediction from the control group. The evaluation metrics include the RMSE and Pearson correlation coefficient (PCC) of the mean expression of each gene between observed treatment group and counterfactual predictions from control group (we compare the mean since the true counterfactual outcome of control cells are not accessible in real data), as well as the maximum mean discrepancy (MMD) between the observed and predicted distributions.

In simulated data, regardless of the accessibility of cell-type labels, scDRP consistently demonstrated robust prediction performance, characterized by significantly lower MMD and RMSE when comparing the predicted and true treated sample distributions, alongside a notably higher PCC (Supplementary Fig.S7). Furthermore, we proceeded to validate the counterfactual prediction performance on 4 real datasets: Kang et al. [14], Dong et al. [6], Hagai, et al. [15], and Norman et al. [16]. These datasets encompass both chemical and genetic perturbations, including experiments on pure cell lines subjected to different perturbations, as well as real tissue cells comprising multiple cell types and even multiple species, providing broad representativeness. As shown in Fig. 2b (known cell-type setting) and Supplementary Fig. S10 (unknown cell-type setting), scDRP exhibits consistently superior performance across nearly all datasets. UMAP visualizations of counterfactual pseudo-treated samples generated from control cells (Fig.2c) further confirm this: the counterfactual distributions produced by our method almost completely overlap with the observed treatment distributions, whereas other methods exhibit substantial gaps.

As shown by the UMAP visualizations (Supplementary Note SN5, Fig. S5-S6), the learned latent spaces exhibit clear disentanglement: *z*_*u*_ aligns stimulated and control cells within distinct cell-type clusters, while *z*_*d*_ resolves treatment-specific variations, even without explicit cell-type labels. This precise separation of features underpins the accuracy of our subsequent conditional OT predictions. The selection of hyperparameters is also a subtle issue. In real-world data, when explicit cell-type labels are used, it is better to impose no or a weaker additional condition on ***z***_*u*_ (Supplementary Fig. S9), while without known explicit cell-type labels, applying a moderate conditional often achieves better results (Supplementary Fig. S11). It is also worth noting that, purely matching by confounding (***z***_*u*_, i.e., setting *β* = 0), similar to the approach adopted by CINEMA-OT, showed very poor results.

### 2.4 Improved ITE estimation and biclustering of ITEs enable finer-grained capture of heterogeneous response patterns

To demonstrate the behavior of our approach, which features more accurate ITE estimation via non-linear disentanglement and conditional OT, alongside biclustering-based (rather than uni-dimensional clustering) downstream analysis for elucidating the biological heterogeneity of perturbation responses, we analyzed the same scRNA-seq dataset of rhinovirus (RV) and cigarette-smoke extract (CSE) exposure in primary human bronchial organoids as CINEMA-OT [6]. The UMAP visualization demonstrates that our more accurate ITE estimates better preserve the heterogeneity in responses to exposure across different cells (Fig. 3a).

**Fig. 3.**
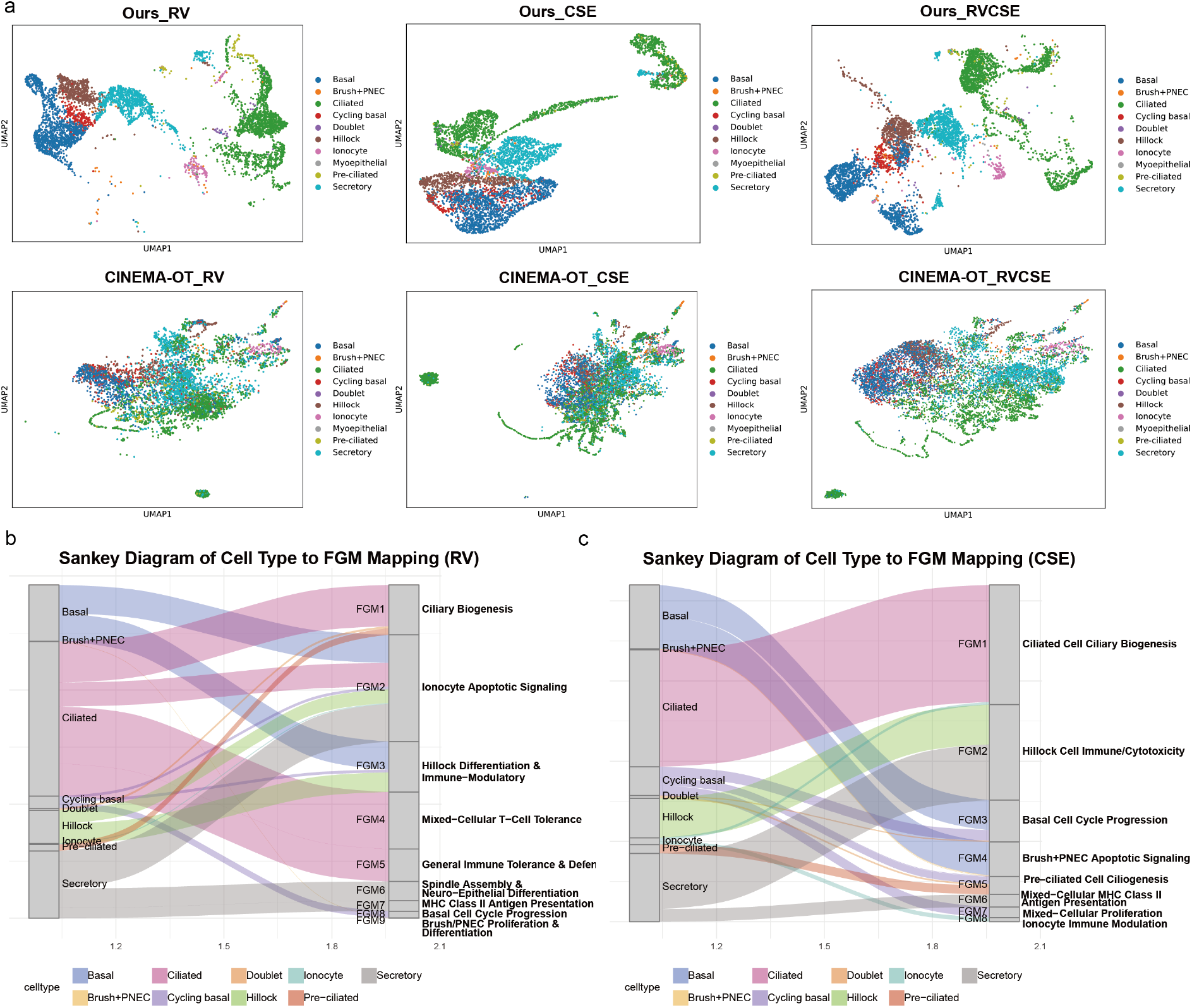
Biclustering of improved ITEs reveals fine-grained heterogeneous response patterns under RV and CSE exposures. **a**. UMAP visualization of the ITE matrix, clustering cell subgroups by their response profiles across different exposures. **b**. Sankey diagram illustrating the enrichment of FGMs within specific cell types following RV exposure. **c**. Sankey diagram illustrating the enrichment of FGMs within specific cell types following CSE exposure.

Our analysis corroborates previous findings of CINEMA-OT [6] that RV treatment induces heterogeneous, cell-type-specific effects, including the activation of apoptotic pathways and general inflammatory responses. Our more accurate ITE estimation and biclustering approach, however, resolves these effects with greater granularity by identifying distinct functional gene modules (FGMs) (Fig. 3b, Supplementary Table S2, Fig. S14). For instance, we specifically localize the intrinsic apoptotic signaling pathway to the Ionocyte-dominated FGM2 module and identify an immune-related transcriptional program enriched for genes annotated to the natural killer cell-mediated cytotoxicity pathway within the Hillock-specific FGM3 module.

More importantly, our analysis reveals major biological programs that were not highlighted in the previous computational analysis. While the prior study prominently highlighted a type I interferon signature, our ITE biclustering strategy uniquely uncovers two dominant biological programs: (1) a strong ciliary biogenesis-related signature (cilium organization, motile cilium assembly) localized to the Pre-ciliated/Doublet FGM1 module, and (2) a strong cell-cycle progression signature (chromosome segregation, spindle organization) enriching in both the Cycling basal (FGM8) and Brush+PNEC (FGM9) cells. These findings suggest that our approach successfully disentangles the RV ITEs into more discrete biological processes, revealing that cell-cycle and ciliogenesis perturbations [17] are central, additional components of the cell-type-specific response that were not resolved by the previous analysis.

Similarly, our analysis confirms that CSE induces a heterogeneous, cell-type-specific response, with notable effects in Basal, Ciliated, and Secretory populations. Moreover, it uncovers a more complex set of biological programs than previously reported (Fig. 3c, Supplementary Table S3, Fig. S15). While prior work highlighted metabolic perturbations, namely Reactive oxygen species pathway, Fatty acid metabolism, and Xenobiotic metabolism, our analysis identifies two dominant, previously unrecognized axes of the CSE ITE.

The first is a widespread disruption of ciliary biogenesis, characterized by the overwhelming enrichment of cilium organization and cilium assembly pathways in both Ciliated (FGM1) and Pre-ciliated (FGM5) cell modules [18]. The second is a strong proliferative response, defined by chromosome segregation and cell cycle G2/M phase transition terms, which we localized to the Basal/Cycling basal compartment (FGM3) [19]. Furthermore, our approach resolves highly discrete immune and stress modules entirely missed by the prior analysis, including a immune-related transcriptional program enriched for genes annotated to the natural killer cell-mediated cytotoxicity pathway in Hillock (FGM2) and Ionocyte (FGM8) cells, and an intrinsic apoptotic transcriptional program within the Brush+PNEC population (FGM4) [20].

### 2.5 Biclustering on estimated ITE matrix reveals cell-type specific immune response to interferon stimulation

We then analyzed the heterogeneous responses of different types of human peripheral blood mononuclear cells (PBMC) to interferon IFN-*β* stimulation [14], which is a potent antiviral and immunomodulatory cytokine that drives a highly coordinated immune program. Visualization of the ITE matrix obtained from scDRP reveals clear clustering of different cell types, demonstrating the heterogeneous effects of IFN-*β* on different immune cells, while the ITE obtained from CINEMA-OT fails to achieve this (Fig. 4a). By further performing biclustering visualization of the ITE matrix obtained from scDRP reveals clear clustering of different cell types, demonstrating the heterogeneous effects of IFN on different immune cells. In contrast, the ITE obtained from CINEMA-OT fails to achieve this. After merging the resulting biclusters [21], we identified eight FGMs.

**Fig. 4.**
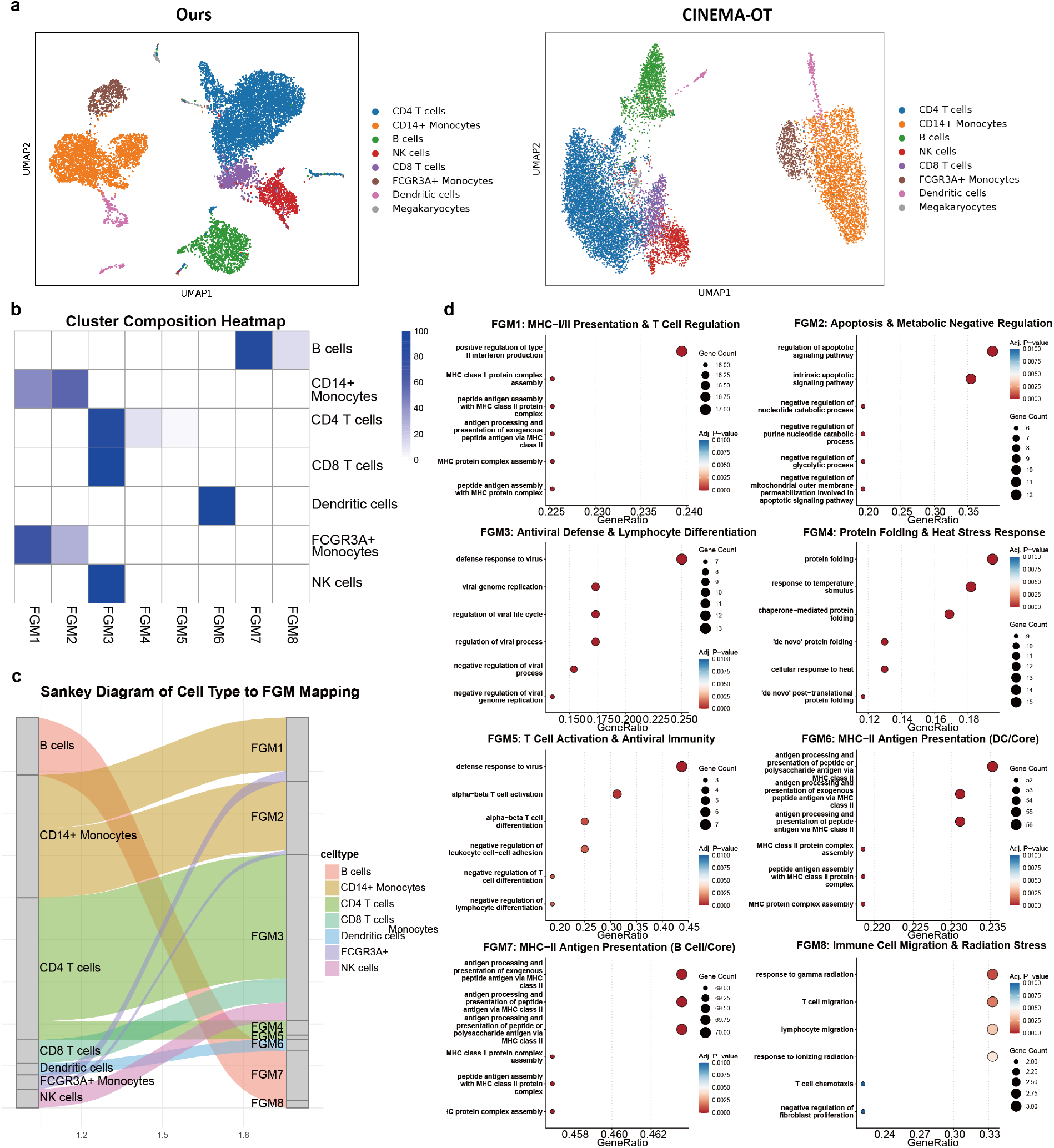
Cell-type specific immune response to interferon stimulation. **a**. UMAP visualization of ITE matrix estimated via scDRP (ours) and CINEMA-OT. **b**. Heatmap on FGM composition among each cell type. **c**. Sankey diagram of cell type to FGM mapping. **d**. GO enrichment analysis reveals functional roles of each FGM. The y-axis displays the enriched GO terms, and the x-axis represents the gene ratio for each term. The size of each dot corresponds to the number of enriched genes in that term, while the color gradient reflects the enrichment *P* -value, scaling from blue (larger *P* -values) to red (smaller *P* -values). The summary on functions of each FGM is shown in the subtitle.

We assigned each cell to the bicluster-derived FGM it belongs to, and then computed the proportion of cells associated with each FGM among each cell type. The results (Fig. 4b-c) reveal distinct patterns associated between cell types and FGMs in response to IFN-*β* stimulation, and we further link them with biological functions through GO enrichment analysis [22] (Fig. 4d, Supplementary Table S4 & Fig. S16). Effector cells, such as CD4^+^ T, CD8^+^ T, and NK cells, all engage FGM3, which relates to antiviral defense and lymphocyte differentiation, confirming the primary role of IFN-*β* in activating the machinery for viral clearance [23]. The CD4^+^ T cells are also associated with FGM4, which is related to functions such as protein folding and endoplasmic reticulum stress. This suggests that the activation of T cells places increased demands on protein synthesis and secretory pathways, accompanied by intracellular homeostatic stress. This heightened metabolic activity is intrinsically linked to cellular homeostatic mechanisms, including the unfolded protein response (UPR) which manages endoplasmic reticulum (ER) stress. It suggests that as T cells ramp up the synthesis and secretion of cytokines and enhance antigen presentation in response to IFN stimulation, they simultaneously engage pathways involved in maintaining proteostasis and alleviating ER stress caused by the high protein load [24].

Meanwhile, antigen-presenting cells (APCs), including DCs (FGM6) and B cells (FGM7), converge on MHC-II Antigen Presentation programs, highlighting a rapid mechanism to potentiate adaptive immunity. Besides, the IFN-*β* stimulus links B cells with FGM8 (Immune Cell Migration) as well, indicating it also activates programs related to cell homing and localization [25]. This suggests that IFN-*β* is consistent with enhanced immune-cell trafficking and migration programs, facilitating essential T-B cell interactions necessary for antibody production.

Most critically, IFN-*β* drives divergent functional fates within the monocyte compartment: FCGR3A^+^ monocytes are skewed toward an activating phenotype, engaging FGM1 (MHC-I/II Presentation and T Cell Regulation), whereas the CD14^+^ monocytes preferentially upregulate FGM2 (Apoptosis and Metabolic Negative Regulation). This functional dichotomy suggests that IFN-*β* employs FGM2 as a crucial negative regulatory mechanism, consistent with increased apoptotic and negative regulatory programs of the CD14^+^ subset to tightly control the magnitude and duration of the immune response.

### 2.6 Chronic myeloid leukemia cells exhibit heterogeneous responses to CRISPR perturbations of different genes

Next, we analyzed the effects of eight different CRISPR-activation conditions (single- and two-gene) on chronic myeloid leukemia (K562) cells [16]. We estimated the ITE of control-group cells under each perturbation (Fig. 5a), performed biclustering of the ITEs for each perturbation separately, and then aggregated the results to obtain seven FGMs.

**Fig. 5.**
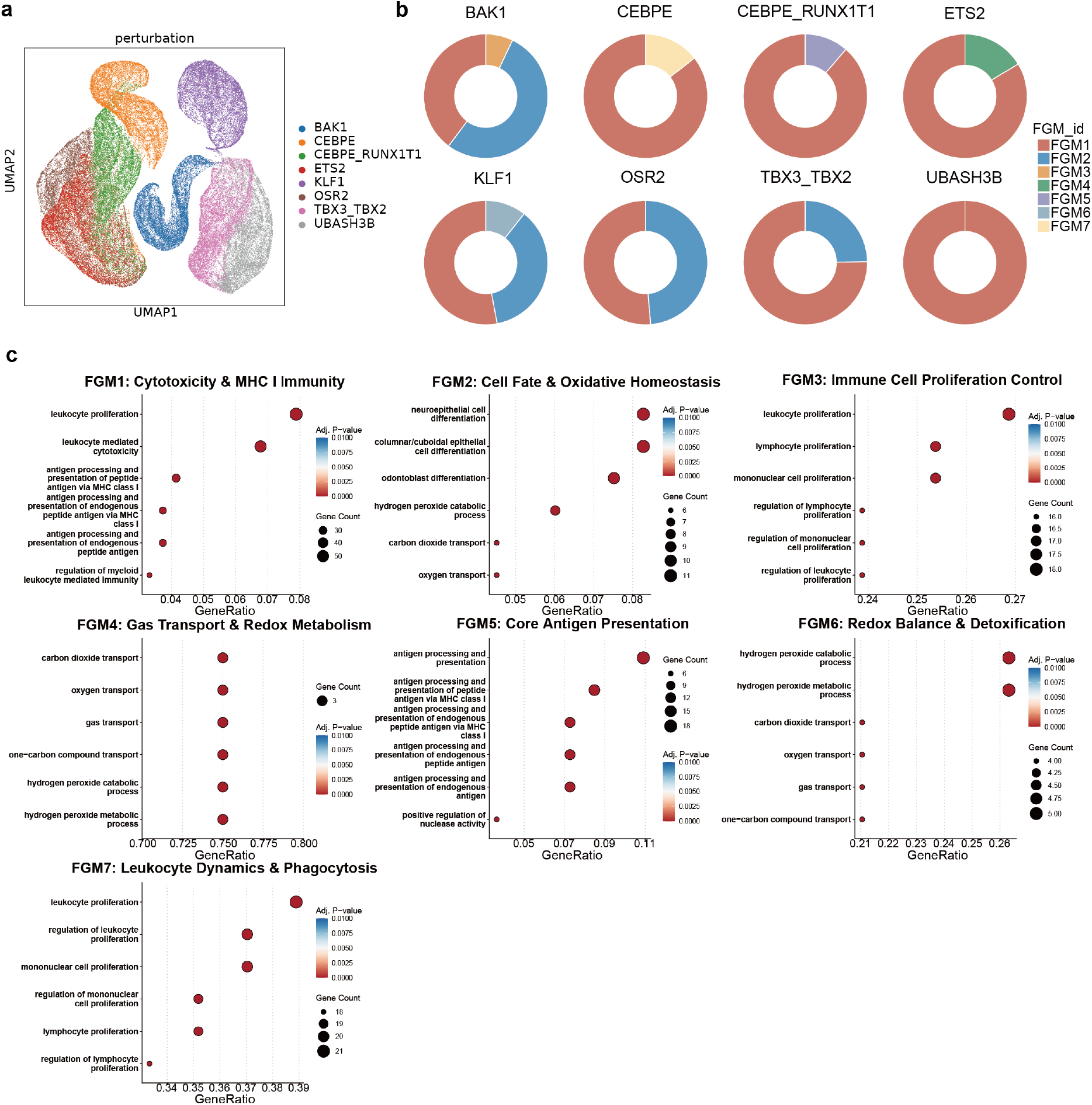
Heterogeneous responses to CRISPR perturbations. **a**. UMAP visualization of the ITE matrix for control cells under different perturbations. **b**. Proportions of FGMs under different perturbations. **c**. GO enrichment analysis reveals functional roles of each FGM. The y-axis displays the enriched GO terms, and the x-axis represents the gene ratio for each term. The size of each dot corresponds to the number of enriched genes in that term, while the color gradient reflects the enrichment *P* - value, scaling from blue (larger *P* -values) to red (smaller *P* -values). The summary on functions of each FGM is shown in the subtitle.

Results (Fig. 5b) show that FGM1 (MHC I antigen presentation and cytotoxicity) is the dominant functional module across almost all eight CRISPR perturbations, comprising 100% of the response for *UBASH3B* and over 75% for several other genes. This strongly confirms that the predominant transcriptional response of K562 cells to genetic perturbation involves antigen-presentation-related, proliferative and apoptotic programs. For example, miR-125b targeting of BAK1 in K562 cells has been shown to suppress apoptosis and promote proliferative signals [26].

However, specific genes show a unique divergence into other critical pathways. FGM2, which relates to cell differentiation and oxidative stress, plays a major role, particularly in the perturbations of *BAK1* (53.1%), *OSR2* (48.7%), and *KLF1* (36.5%). KLF1 has been shown to orchestrate almost all aspects of terminal erythroid differentiation in both human and murine systems [27], and in CML leukemic stem cells, shifts in oxidative stress via mitochondrial metabolism are coupled to differentiation when certain regulatory pathways are inhibited (ULK1, etc.) [28]. This suggests these genes are critical nodes linking cell death signaling to the leukemia cell’s differentiation status and oxidative tolerance, where their activation may promote differentiation-associated and oxidative-stress-related transcriptional programs.

Furthermore, specific transcription factor perturbations reveal distinct functional consequences relevant to the myeloid lineage and immune evasion (Fig. 5c, Supplementary Table S5 & Supplementary Fig. S17). CEBPE’s overexpression, while primarily FGM1, uniquely activates FGM7 (Phagocytosis), reflecting its key role in regulating myeloid functional maturation, especially granulocytic differentiation [29]. The co-perturbation of *CEBPE* and *RUNX1T1* is the only one to engage FGM5 (MHC I Assembly) significantly, suggesting a potential mechanism for modulating antigen-presentation-related pathways in CML cells. Lastly, the unique link of *ETS2* to FGM4 (Gas Transport and Detoxification) suggests it is a regulator associated with oxidative-stress and metabolic-response programs of leukemia cells. This is consistent with broader findings that ROS regulation and metabolic reprogramming are central in myeloid leukemias, with antioxidant pathways linked to survival, cellular adaptation, and therapy resistance [30], identifying a potential vulnerability for combined metabolic and genetic targeting. We further examined whether promoter nucleosome architecture is associated with perturbation responsiveness, using promoter CpG island (CGI) status as an indirect sequence-based proxy for promoter nucleosome organization [31, 32]. This proxy is motivated by the sequence determinants of nucleosome organization: CpG and GC content contribute to sequence-encoded nucleosome depletion tendencies [32], and interpretable sequence models trained on MNase-seq data have identified CpG-related features among the dominant predictors of nucleosome organization, with CCG/CGG periodicity serving as a highly discriminative feature between nucleosomal and inter-nucleosomal DNA [33]. CGI status therefore provides a coarse promoter-level indicator of sequence features that influence nucleosome organization at nucleotide resolution. After stringently controlling for baseline expression levels, we observed that genes with non-CGI promoters exhibited modest but consistently larger transcriptional responses than CGI genes across all eight perturbations (Supplementary Fig. S18). This is consistent with previous reports that non-CGI promoters generally display higher nucleosome occupancy, requiring stimulus-dependent chromatin remodeling for activation, whereas CGI promoters tend to adopt nucleosome-depleted and more permissive chromatin architectures [31]. As CGI status is only an indirect sequence-based proxy for nucleosome occupancy, this association remains suggestive and requires further confirmation with direct chromatin-level measurements in future work.

### 2.7 scDRP predicts dose-sensitive cellular response to chemical perturbations by interpolating on latent space

To assess the model’s capacity of generalizing to unseen conditions, we examined cellular responses across a range of chemical perturbation doses. We analysed the sciplex4 dataset [2], which profiles mammary epithelial and alveolar basal epithelial cells treated with seven compounds at multiple doses. To capture dose-specific effects, we extended the perturbation index ***a*** with an additional dimension encoding dose values *v*. For doses with observed cells, we inferred the effects by applying optimal transport in the disentangled latent space, a procedure we term Estimation. For unobserved doses, we employ conditional Gaussian process regression to impute the mean and variance of ***z***_*d*_, for unobserved dose values, and apply residual correction to ensure quantile alignment of samples before and after perturbation while keeping ***z***_*u*_ unchanged. The VAE decoder is then used to generate counterfactual samples for causal effect estimation, which is a strategy we term Extrapolation (see Methods for details).

We estimated the effects of seven chemical perturbations on two cell types in the SciPlex4 dataset using two different methods. For each cell type, we selected the five genes most strongly affected and plotted the CATE–dosage curves (Fig. 6a). The interpolated curves closely overlapped with those estimated at observed dose values, demonstrating the strong generalization ability of our method. In contrast, direct estimates at observed doses exhibited greater variability, as they contained more noise, compared to the smoothed interpolation results obtained from the Gaussian process.

**Fig. 6.**
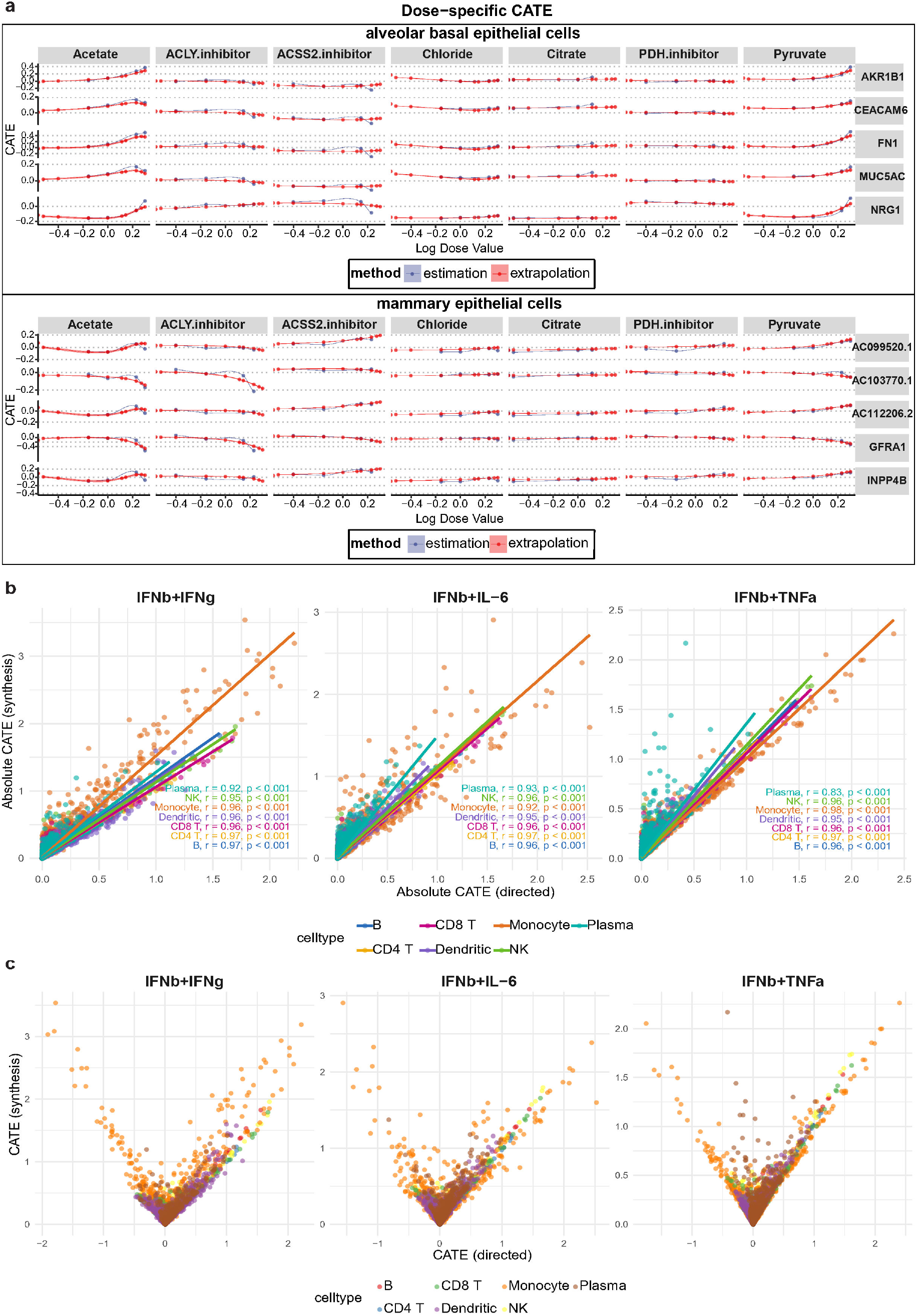
scDRP generalizes to unseen dosage values and treatment combinations. **a**. Dose-response (CATE) curves estimated via estimation (observed dosages) and extrapolation (latent interpolation) for two cell types and seven perturbations in sciplex4 dataset. **b**. Absolute CATE of combined treatments obtained via direct estimation vs synthetic inference from individual treatments for top 3000 genes. The lines are fitted via linear regression for each cell type, with Pearson correlation *r* and *P* -values reported. **c**. CATE of combined treatments obtained via direct estimation vs synthetic inference from individual treatments for top 3000 genes.

We can obtain more detailed information about perturbation effects from the dose–response curves. For example, in alveolar basal epithelial cells, Acetate stimulation leads to a dose-dependent upregulation of *AKR1B1* expression, which may be due to enhanced flux through the glucose-derived polyol pathway under high acetate conditions, increased production of NADPH to buffer oxidative stress, or activation of metabolic reprogramming that favors enzymes like *AKR1B1* [34]. Indeed, *AKR1B1* is known to mediate metabolic reprogramming and enhance stress tolerance in contexts of hyperglycemia and increased energy demand [35].

In contrast, in mammary epithelial cells, stimulation with an ACLY inhibitor exerts a sustained suppressive effect on GFRA1 expression, likely because inhibition of ACLY reduces cytosolic acetyl-CoA supply, thereby limiting lipid and protein acetylation needed for growth factor signaling and transcriptional activation; additionally, decreased ACLY activity can impair metabolic pipelines (e.g. lipogenesis, histone acetylation) [36], crucial for maintaining GFRA1 expression.

### 2.8 scDRP helps infer synthetic effects of multiple treatments via vector composition on ITE space

We further demonstrated scDPR’s extrapolative ability for inferring the synthetic effects of multiple perturbations given observations of single perturbations. We conceptualized the problem as a vector composition issue (see Methods). We validated this approach using a dataset [6] of human PBMCs subjected to single and combined stimulations, including IFN*β* + TNF*α*, IFN*β* + IFN*γ*, and IFN*β* + IL-6. We compared the absolute CATE of each gene within each cell type using ITEs estimated directly from samples subjected to combined stimulations with those calculated by vector addition of the ITEs from each single stimulation.

As shown in the Fig. 6b, one can observe a very strong linear correlation, which demonstrates the effectiveness of our method in inferring combinatorial effects. At the same time, note that our method can only estimate the absolute magnitude of the combined effect, but cannot determine whether the final effect is an upregulation or downregulation of gene expression (as clearly illustrated by the absolute value function effect in the Fig. 6c).

## 3 Discussions

Disentangled representation learning has recently received widespread attention to address the unpaired cross-condition alignment challenge arising from the destructive nature of single-cell measurements [5, 37]. In this study, we introduce scDRP, a framework that enables the estimation of ITEs and counterfactual responses from single-cell perturbation data. By combining disentangled representation learning with conditional optimal transport, scDRP provides a principled way to isolate the true causal impact of perturbations from confounding variation. Our results demonstrate that this formulation not only improves the accuracy of counterfactual prediction but also facilitates interpretable biological insights by linking latent dimensions to functional gene modules. Latent-space interpolation and ITE vector composition illustrate the model’s capacity to generalize beyond observed experimental conditions, positioning scDRP as a versatile computational framework for identifying perturbation effects in high-dimensional single-cell systems.

We address the fundamental problem of causal inference that only one potential outcome is observable for any given cell-through latent space decoupling and counterfactual matching. Given the complexity of biological systems and substantial technical noise, traditional linear models carry a high risk of model misspecification [38]. We therefore opted for a flexible neural network-based approach, which allows us to effectively perform the decoupling while accommodating technical noise and batch effects. We achieve the disentanglement based on the assumption that, given proxies of confounders (such as cell type), the perturbation-dependent and perturbation-invariant latent variables are independent. When cell-type information is unavailable, only partial disentanglement can be achieved. However, by incorporating the distance in ***z***_*u*_ (which encodes confounding information) into the cost function of OT as a soft conditional term, the model retains considerable robustness even in the absence of explicit cell-type labels.

While OT has recently been widely applied in single-cell analysis with unpaired data across different omics modalities, conditions, or time points [6, 39], existing methods are conducted in an intuitive and heuristic manner, lacking explicit underlying assumptions and rigorous theoretical guarantees. In this work, we establish rigorous theoretical assumptions for utilizing conditional OT to perform counterfactual matching. The core idea of counterfactual matching in the latent space stems from the principle that, after controlling for confounders, a treatment may shift the overall population distribution but should not alter the relative position of each individual within the population, that is, the exogenous noise is independent of the treatment, which is commonly adopted in counterfactual inference. In fact, it corresponds to the FréchetHoeffding upper bound on the joint distribution induced by the conditional marginal distributions of potential outcomes after controlling for confounders, specifying the most conservative coupling (i.e., the minimal change assumption) that minimizes the individual causal effects [13]. We implement a soft-constrained conditional OT to jointly perform confounder control (***z***_*u*_) and quantile matching (***z***_*d*_), inspired by the equivalence between one-dimensional OT under convex loss and quantile mapping [7]. Exploring alternative estimation strategies, such as independent residual correction or coupled per-dimension quantile matching, represents an important avenue for future work. Future extensions may also incorporate temporal or spatial modalities to further disentangle dynamic regulatory programs and cellular interactions.

The interpretable product of scDRP is the set of functional gene modules obtained by biclustering the ITE matrix: each module maps directly onto annotated biological processes through enrichment analysis (Figs. 3–5; Tables S2–S5), providing a cell-type-resolved account of which programs change under a perturbation. The disentangled latent representations, serving as an intermediate product to extract and decouple compressed information from high-dimensional and noisy transcriptomic profiles, are deliberately not assigned component-wise individual biological meanings. Although by the identifiability theory underlying our model (Supplementary Note SN1), the sparsity-constrained, independent latent components are recoverable up to some invertible transformation under proper regularity conditions, interpretation is more intuitive and reliable at the level of functional modules and effects rather than latent components.

The study still has several limitations. It rests on identifying assumptions that, as is typical in causal inference, cannot be verified from observational data alone. The disentanglement assumes that response and identity factors are conditionally independent given confounders (or their surrogates). When no confounder proxy is available, only partial disentanglement is achievable, and the soft conditional term in the OT cost mitigates but does not eliminate residual confounding. The counterfactual matching assumes rank preservation within homogeneous subpopulations, a conservative, minimal-change coupling, which may not hold if a perturbation reorders cells’ relative sensitivities. The combinatorial-effect inference recovers only the magnitude of a joint effect and not its direction. Finally, current scDRP models transcription-level response (scRNA-Perturb-seq) only and does not incorporate more sequence- or chromatin-level information. Linking gene-level ITEs to regulatory or sequence features, and extending the framework to perturbation profiling of other modalities such as scATAC and multiome Perturb-seq [40], are important directions for future studies. We believe that as more advanced perturbation sequencing technologies are developed and more comprehensive datasets are accumulated across diverse cell and tissue types with richer perturbations, scDRP will serve as a powerful computational tool. It will help elucidate the cellular and molecular mechanisms in biological systems under perturbations at an unprecedented resolution and facilitate the discovery of novel drug targets.

## 4 Methods

### Basic idea and intuition

Quantifying how a perturbation acts on an individual cell is hindered by the fundamental problem of causal inference: each cell is observed in only one condition, and hence its counterfactual is never directly measured. This is compounded by single-cell heterogeneity, in which a cell’s response is entangled with its intrinsic identity (cell type, state, and other covariates). The central idea of scDRP is to disentangle these two sources of variation in a learned latent space: an identity representation ***z***_*u*_ capturing the perturbation-invariant intrinsic state of a cell, and a response representation ***z***_*d*_ capturing the perturbation-induced change, with an explicit independence constraint enforcing the separation. With identity and response separated, scDRP estimates each control cell’s counterfactual by matching it to perturbed cells of similar identity, so that the two differ only in the perturbation.

The premise behind this matching is that a perturbation acts similarly on cells of similar intrinsic state: within a subgroup of comparable identity, it shifts the cells’ responses while preserving their relative ordering. Under this conditional rank-preservation, a control cell’s counterfactual is the perturbed cell occupying the same response rank within its identity-similar neighborhood. scDRP obtains such matchings by solving a convex optimal-transport problem on the response representation ***z***_*d*_, with a soft penalty on ***z***_*u*_ dissimilarity that conditions the coupling on intrinsic identity. The individual treatment effect is then read off as the difference between each cell’s counterfactual and its observed state.

### 4.1 Disentangled representations with minimal-change principle

Suppose we have the observed gene expression ***x***, perturbation index ***a***, and cell type index ***w*** (optional), as well as some technical covariates like sequencing batch index ***c*** (optional). We assume observed ***x*** ∈ ℝ^*p*^ is generated from latent gene programs ***z*** ∈ ℝ^*d*^ (*d* ≪ *p*) and that only a subset of ***z*** is affected by the perturbation. Under the assumption that ***z***_*d*_ are ***z***_*u*_ are conditionally independent given confounding ***w***, we employ the following VAE architecture to learn the disentangled representations of gene expressions.

- Generative network: We assume the generating process: *p*(***x***) = *p*(***z***)*p*(***c***)*p*(***x***|***z, c***), where 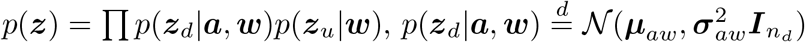 and 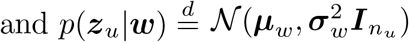. There are flexible choices to specify *p*(***x***|***z, c***) and reconstruction loss (log likelihood) accordingly. For example, (1) if the log-normalized data are provided as ***x***, we can use a half Gaussian distribution |*N*(*f* (***z, c***), Σ)| and use mean squared error (MSE) loss as the reconstruction loss; (2) if the raw count data are provided, we may assume that *p*(***x***|***z, c***) follows a zero-inflation negative binomial (ZINB) model [41] 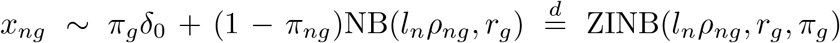, where *l*_*n*_ is the library size of cell *n, ρ*_*ng*_ is the mean expression proportion of gene *g* in cell *n* generated by some injective functions *ρ*_*ng*_ = *f* (***z, c***), *r*_*g*_ is the dispersion factor of gene *g, π*_*ng*_ is the dropout rate of gene *g* in cell *n*, and we have ℒ_recon_ = −∑_*n*_ log {*P*_ZINB_(***z***_*n*_; *l*_*n*_***ρ***_*n*_, ***r, π***)}.
- Inference network: We use the following inference network to approximate the generative process: with encoders *q*(***z***_*d*_|***x, a, w***) and *q*(***z***_*u*_|***x, w***), and we have

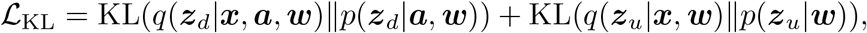

where KL denotes the Kullback-Leibler divergence.

To encourage disentanglement, we use *β*-VAE (which introduces a tunable hyperparameter *β* on the KL term) [42] and impose an additional independence loss [43] on VAE ELBO by Hilbert-Schmidt conditional Independence Criterion (HSCIC) [9, 44] to impose ***z***_*d*_ ⊥ ***z***_*u*_|***w***, i.e., ∑_ind_ = HSCIC(***z***_*d*_, ***z***_*u*_ | ***w***), where

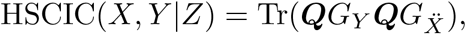

which measures conditional dependence through the projection operator ***Q*** = ***I*** − ***V*** with ***V*** = (***G***_*Z*_ + *nϵ****I***)^−1^***G***_*Z*_, multiplied with the centered Gram matrix with the extended variable kernel 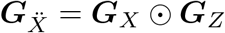.

To encourage identifiability via sparsity constraints [45] and automatically select the optimal latent dimensions of ***z***_*d*_ and ***z***_*u*_, we introduce a gate variable ***m*** on ***z*** (i.e., use ***m*** ⊙ ***z*** as the real latent factors for generating process) and then imposing hard concrete distribution on *m* and shrinkage the latent dimension by 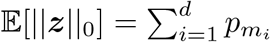, where 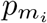 is the activation probability of each gate calculate via the hard concrete distribution [10].

The final loss is hence

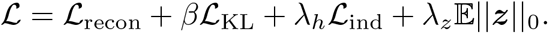

### 4.2 Individual effect estimation and counterfactual prediction

In our generative model framework, the counterfactual outcomes for control group cells after treatment can be generated using the decoupled latent representations. Suppose we obtain the disentangled 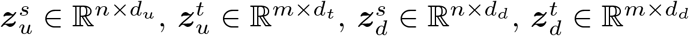, the perturbation-invariant latent components ***z***_*u*_ will remain unchanged, while the key issue is to transfer ***z***_*d*_ for samples observed in the control domain to the treatment domain. To achieve this, we transfer ***z***_*d,i*_ from source distribution (in control group) 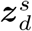 to the target distribution (in the corresponding treatment group) 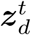 while conditioning on similar *z*_*u,i*_. We resort to conditional rank-preserving assumption, which gives a conservative estimate that minimizes the overall ITE (minimal change principle), corresponding to the Fréchet-Hoeffding upper bound on conditional coupling 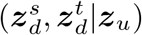 [13].

Ideally, it can be achieved by defining the cost as

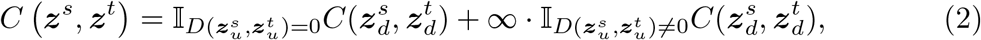

to constraint the shift of ***z***_*d*_ distribution under the condition of the same ***z***_*u*_. However, in reality, it would be extremely hard or impossible to find samples in two domains that are exactly matched. Even if we have some class labels as surrogates (e.g. cell types), we can do conditional transfer within each class, but the heterogeneity within each class (e.g. subclass) is still neglected. And therefore, we relax to

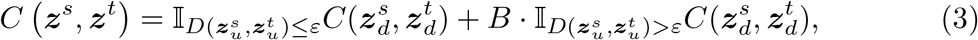

where *B >* 0, which is equivalent to the following cost function

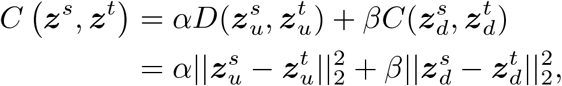

where *α* controls the strength of conditionals and *β* controls the weights of transport cost, and we can get the optimal coupling matrix via optimal transport (OT) solvers

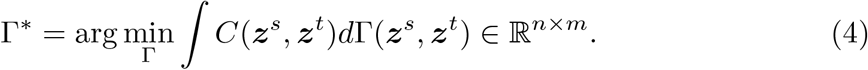

We take both distance measure 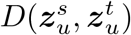 and transport cost 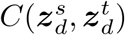 as squared Euclidean distances. The rationale is twofold. First, the strict convexity of this cost ensures that in the one-dimensional case, optimal transport is equivalent to quantile matching [7], while in the multi-dimensional case it can be viewed as the sum of costs across each dimension. This makes it conceptually consistent with our assumption that perturbations correspond to rank-preserving shifts along each latent dimension. Second, since we perform disentanglement using a VAE, where the KL prior encourages the latent space to approximate a low-dimensional Gaussian distribution, adopting squared Euclidean distance as both the distance metric and the transport cost in the latent space is natural and effective.

When *α* = 1 and *β* = 0, the procedure resembles a smoothed version of *k*NN, where matching between pre- and post-perturbation cells relies solely on the invariant component *z*_*u*_. In contrast, when *α* = 0 and *β* = 1, the procedure amounts to applying no conditional control, directly transferring the distribution of *z*_*d*_ from the control group to the treatment group in a quantile-preserving manner [7]. In practice, when cell types are known, we first perform domain adaptation in the latent space within each cell-type group, in which case a relatively small value of *α* can be used. By contrast, when cell types are unknown, domain adaptation is applied jointly across all samples, and a larger value of *α* is required.

Upon obtaining the optimal coupling matrix Γ, we induce distributional shift through barycentric projection. We consider two alternative strategies: (i) performing barycentric projection over the entire latent space,

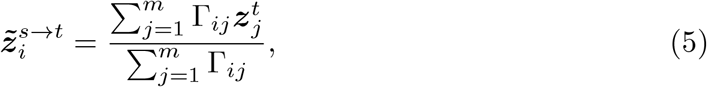

and (ii) applying barycentric projection to ***z***_*d*_ only while keeping the original ***z***_*u*_ unchanged.

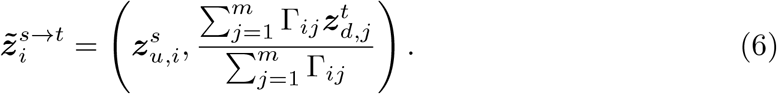

We can then generate counterfactual samples for control group cells after treatment as 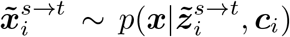, ***c***_*i*_), and the individual treatment effect (ITE) for the control cells when being treated can be estimated as

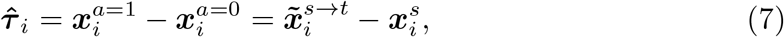

and the conditional average treatment effect (CATE), or more precisely conditional average treatment effect of the untreated (CATU), within a specific cell type *q* is 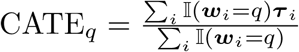.

### 4.3 Biclustering on the ITE matrix

The ITE matrix contains information about the heterogeneity of responses to the same perturbation across different cells and genes. To uncover this heterogeneity, we employed the BCPlaid algorithm [46] (available via R package biclust, with the default parameters) to perform biclustering analysis on the estimated ITE matrix, aiming to identify subpopulations of cells and gene sets that exhibit specific response patterns.

We first perform biclustering on cell groups under different conditions (e.g., distinct cell types or perturbations), and then merge and summarize the resulting biclusters. During summarization, we first remove genes that are broadly present across all biclusters within a given condition, as these typically reflect housekeeping or background signals rather than condition-specific differences. Next, we merge biclusters within each condition, followed by merging biclusters across different conditions [21].

We merge biclusters within the same condition using the overlap score (OS) to pay more attention on exclusion, defined as

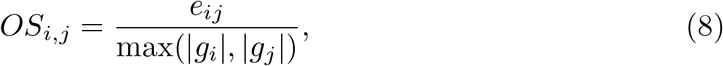

where *e*_*ij*_ denotes the number of genes only appears in biclusters *i* and *j*, and *g*_*i*_ represents the set of genes in bicluster *i*. Bicluster pairs with *OS >* 0.03 are combined. We subsequently merge them by emphasizing the degree of overlap since biclusterings across different conditions are independent, quantified using the “intersection-over-union” (IoU) criterion

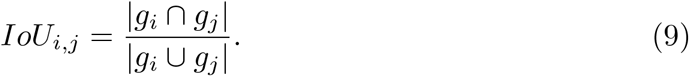

Pairs of biclusters with *IoU >* 0.3 are combined. Through this two-step merging procedure, we ultimately obtain functional gene modules (FGMs) that capture condition-specific biological signals.

Cells are subsequently assigned to FGMs based on their bicluster membership, allowing us to characterize changes in FGM proportions across different conditions. Furthermore, we apply the R package clusterProfiler [22] to perform enrichment analysis on these differential FGMs, thereby linking the identified modules to specific functional pathways and biological processes.

### 4.4 Unseen dose-specific responses predicted via latent space interpolation

To quantitatively describe how cells response to a particular treatment with covariates such as chemical dose values (*v*, typically log-transformed), we concatenate the perturbation dose values as one component with the one-hot encoded perturbation index to form a new perturbation index (*a*), which is then input into the model. For the dose values of cells in the observational dataset that have been subjected to the corresponding perturbation (*v*_obs_), we can directly apply the previously mentioned method to estimate the effect size of that dose treatment on the cells (The process we refer to as Estimation).

In contrast, for dose values that have never been observed in the dataset (*v*_unseen_), assuming ***z***_*u*_ remaining unchanged, we interpolate the unknown ***z***_*d*_(*v*_unseen_) by using Gaussian Process (GP) regression (conditioning on ***w***) to predict the dose-dependent mean 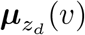 (*v*) and standard deviation 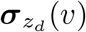 (*v*), with hyperparameters optimized by maximizing the log marginal likelihood. Personalized counterfactuals were generated based on a residual preservation principle. For each cell in the control condition, we calculated its standardized latent residual, 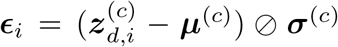, which captures its unique deviation from the control population. Assuming this residual is conserved, the counterfactual latent state for a new dose *v*_new_ was synthesized by transplanting this residual onto the predicted distribution: 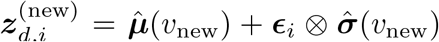. The final high-dimensional cell profile was then generated by passing the complete counterfactual vector, 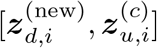, through the pre-trained VAE decoder (The process we refer to as Extrapolation).

### 4.5 Synthetic effect estimation of multiple treatments through ITE vector composition

To infer the combined effects of multiple perturbations from a single-cell dataset that includes responses to different individual perturbations, we analogize the superposition of different perturbations to the addition of vectors (Fig. 1d). Through decoupled learning, we can estimate the ITE for each cell *i* in the control group after receiving perturbation 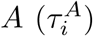 and perturbation 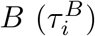 separately. We can then calculate the correlation coefficient *r*_*AB*_ between *τ* ^*A*^ and *τ* ^*B*^ among all cells, which can be interpreted as the cosine of the angle between the vectors representing the effects of these two treatments. Subsequently, using the parallelogram law of vector addition and the cosine theorem, we can infer the absolute synthetic effect 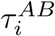 when the cells simultaneously receive treatments *A* and *B* as

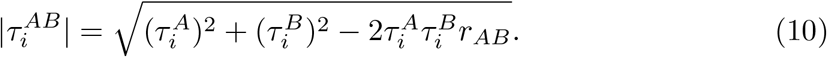

Given that angles summing to 360 degrees share an identical cosine, we can only estimate the magnitude of the combined effect, while its direction (up- or down-regulation) cannot be discerned (Fig. 1d), shown in the V-shape plot (Fig. 6c).

### 4.6 Simulations

We simulate single-cell perturbation dataset with the following procedures. We first simulate *N* cells in the control group, where the latent components are generated from Gaussian mixture model 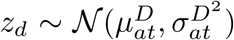 and 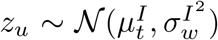 with cell type labels *t* sampled from a Dirichlet distribution *Dir*(*K*). Then, for treatment group, we use the same *z*_*u*_ and transform *z*_*d*_ to 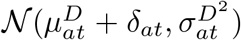, where *δ*_*at*_ ∼ *N* (1, 0.5^2^), and ensure that the ranks of the corresponding samples remain consistent before and after the shift. The mean expression of gene *g* for each sample *µ*_*ng*_ is generated as 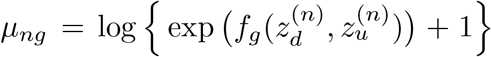, where *f*_g_ (·) is a non-linear neural network, where we impose selective masks on latent components when generating different genes to increase the sparsity. The dispersion factors *r*_*g*_ and dropout rates *π*_*g*_ are sampled to make highly expressed genes have lower Biological Coefficient of Variation (BCV, here we specify dispersion factor *r* = 1*/α* = 1*/*BCV^2^) and lower dropout rate, i.e., 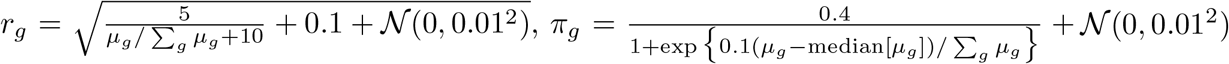. The gene expressions are then sampled from zero-inflated negative binomial distributions *x*_*ng*_ ∼ *π*_*g*_*δ*_0_ + (1 ™ *π*_*g*_)NegBin(*µ*_*ng*_, *r*_*g*_). We randomly sample *N/*2 cells in control and *N/*2 cells in treatment groups to get the final observed single-cell dataset.

### 4.7 Evaluations

For the simulated data, we computed the RMSE of the estimated ITE and CATE on all genes. For the real datasets, we compared the observed treatment cells with the counterfactually generated “pseudo-treatment” samples from the control cells. We evaluated the RMSE and PCC of their mean gene expression levels within top 3000 HVGs as we cannot access the individual-level counterfactual truth. We also evaluated the MMD between the observed treatment 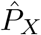 and counterfactually predicted distributions 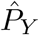. To avoid spurious dilution of MMD values caused by uniformly (un)expressed background or housekeeper genes, MMD is calculated using top 200 HVGs by the V-statistic (for convenience in vectorized implementation) [47],

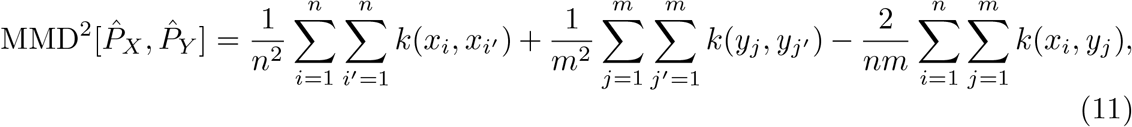

where *k*(·, ·) is the Gaussian (Radial Basis Function, RBF) kernel 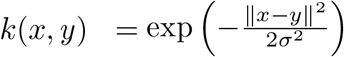. The bandwidth *σ* is chosen using the median heuristic, where it is set equal to the median of the Euclidean distances between all pairs of samples in the dataset. We also visualized the results using UMAP.

## Supplementary information

Supplementary materials are available online.

## Acknowledgements

We sincerely appreciate Dr Richard Border and Dr. Martin Jinye Zhang for their valuable comments and suggestions on this work. Fig. 1 is created with BioRender (https://BioRender.com).

## Key Points

- We overcome the destructive nature of single-cell Perturb-seq by locating a control cell’s counterfactual twin as its treatment-group neighbor that shares a similar intrinsic identity while preserving its relative rank along the perturbation response axis.
- scDRP employs a sparsity-regularized beta-VAE to disentangle invariant intrinsic identity from perturbation-induced response, with conditional optimal transport to estimate individualized treatment effects (ITEs) from unpaired populations.
- scDRP accurately predicts individual counterfactuals and robustly generalizes to unseen doses and combinatorial treatments via latent space interpolation.
- Validated across simulations and real datasets, scDRP uncovers cell-type-specific heterogeneous responses, revealing novel ciliary biogenesis and cell-cycle signatures.

## Declarations

### Ethics approval and consent to participate

Not applicable.

### Consent for publication

Not applicable.

### Data availability

The Kang and Hagai datasets are available via pertpy package (http://pertpy.readthedocs.io/en/latest/index.html). The Norman and sciplex4 datasets are available via scPerturb (https://zenodo.org/records/13350497). The Dong dataste are available from DRYAD (https://datadryad.org/dataset/doi:10.5061/dryad.4xgxd25g1).

### Code availability

Codes of scDRP are available at https://github.com/sjl-sjtu/scDRP. CINEMA-OT and scGen are implemented with Python library pertpy (http://pertpy.readthedocs.io/en/latest/index.html).

### Author contribution

J.S., P.S. & K.Z. conceptualize the methods; J.S. implemented the methods, conducted formal analysis and wrote the original manuscript; P.S. helped interpreting the results; K.Z. supervised the study; J.S., P.S., & K.Z. revised the final manuscript.

## Supplementary Materials

### Supplementary Notes

#### SN1 Identifiability theory of non-linear disentanglement

##### 1.1 General Theory

Recent advances in the theory of disentangled and nonlinear ICA models have progressively relaxed the conditions required for identifiability, i.e., the ability to recover unique latent variables up to subgroup or component-wise invertible transformations and permutations.

Zheng et al. [1] first established Structural Sparsity as a sufficient condition for identifiability in fully unsupervised nonlinear ICA, showing that when each latent factor uniquely determines at least one subset of observed variables (i.e., the intersection of their parents identifies that source), the true latent components can be recovered without any auxiliary variables.

Building upon this, Zheng and Zhang [2] generalized the theory to more realistic settings by introducing partial sparsity, undercompleteness (more observed than latent variables), and partial source dependence. They proved that if structural sparsity and independence hold for only a subset of sources, those sources remain component-wise identifiable, while the remaining (possibly dependent) ones are identifiable up to a subspace. The framework also allows auxiliary variables with only a small number of distinct values to assist identifiability, extending nonlinear ICA to mixed or grouped latent structures.

Most recently, Li et al. [3] unified two previously separate lines of identifiability research, i.e., sufficient distributional changes across auxiliary variables and sparse mixing structures, into a single complementary framework. They demonstrated that sparsity in the mixing process can compensate for insufficient domain variation, while diverse conditional distributions can mitigate violations of sparsity. Under smoothness, conditional independence, and either moderate distributional changes or partial sparsity, both subspace- and componentwise identifiability can be achieved with milder assumptions than required by classical nonlinear ICA results.

These works broaden the identifiable regimes for disentangled representation learning, where the theoretical insights are particularly relevant to single-cell generative modeling, where identifying biologically meaningful latent factors, such as cell identity or state, is paramount. In practice, the structural sparsity corresponds to the modular nature of gene regulatory networks, where each latent factor (e.g., a specific transcription factor) regulates only a sparse subset of marker genes. Simultaneously, the auxiliary variables manifest as experimental metadata, such as perturbation conditions (e.g., drug treatment or gene knockdown). By leveraging the sparse connectivity of gene modules alongside the distributional shifts induced by these perturbations, generative models can achieve robust identifiability, effectively disentangling intrinsic cellular programs from extrinsic experimental effects.

##### 1.2 Identifiability under noisy observations

While the aforementioned theories primarily focus on deterministic or noiseless mixing processes, real-world generative mechanisms are inevitably corrupted by observation noise, which fundamentally alters the identifiability landscape. Khemakhem et al. [4] provided the rigorous identifiability results for deep latent-variable models under additive noise by embedding nonlinear ICA theory into the variational autoencoder framework. Specifically, for a noisy generative process of the form *x* = *f*(*z*) + *ε*, where *f*is an injective nonlinear function and *ε* denotes independent noise, we introduced a conditionally factorized latent prior *p*(*z* | *u*) whose parameters vary with an auxiliary observed variable *u*. By modeling each latent component with an exponential family whose sufficient statistics are modulated by *u*, the joint distribution *p*(*x, z* | *u*) becomes identifiable up to component-wise invertible transformations and permutations, even in the presence of noise and under undercomplete settings. This result demonstrates that distributional variation across auxiliary variables can overcome the fundamental non-identifiability induced by nonlinear mixing and observation noise, extending classical nonlinear ICA identifiability to realistic stochastic generative models. Importantly, their theory establishes consistency of VAE-based maximum likelihood estimation, showing that both latent variables and generative mechanisms can be recovered (up to trivial indeterminacies) rather than merely matching the marginal data distribution.

Classical structural sparsity-based identifiability results enforce sparsity in the Jacobian of the mixing function to restrict the dependency graph between latent and observed variables. However, explicitly computing and regularizing the Jacobian becomes prohibitively expensive in deep generative models. Instead, we adopt a complementary surrogate by constraining the effective latent dimensionality through learnable gating variables with *ℓ*_0_ regularization. This mechanism encourages only a small subset of latent components to actively participate in the generative process, thereby implicitly inducing sparse dependency structures between latent and observed variables. From a functional perspective, restricting latent participation reduces the expressiveness of admissible mixing functions and limits the class of nonlinear reparameterizations responsible for non-identifiability. While not equivalent to explicit Jacobian sparsity, this latent selection strategy serves as a practical approximation that promotes structural sparsity in deep models with significantly lower computational overhead.

##### 1.3 Regularization on sparsity

Classical structural sparsity-based identifiability results enforce sparsity in the Jacobian of the mixing function to restrict the dependency graph between latent and observed variables. However, explicitly computing and regularizing the Jacobian becomes prohibitively expensive in deep generative models. Instead, we adopt a complementary surrogate by constraining the effective latent dimensionality through learnable gating mechanism 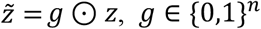 with *ℓ*_0_ regularization [5]. This mechanism encourages only a small subset of latent components to actively participate in the generative process, thereby implicitly inducing sparse dependency structures between latent and observed variables. Moreover, by encouraging only a small number of active gates, the model automatically infers the effective latent dimensionality, mitigating the overparameterization that otherwise exacerbates non-identifiability in nonlinear generative models. Though slightly stronger than classical entry-wise Jacobian sparsity, this latent selection strategy serves as a commonly used and practical approximation that promotes structural sparsity in deep models with significantly lower computational overhead [6].

#### SN2 Conditional optimal transport on disentangled latent space

Recall that we assume that the observed variables ***x*** are generated by both domain (perturbation)-dependent latent factors ***z***_*d*_ ∼ *p*_*d*_(***z***_*d*_|***a, w***) and domain-invariant latent factors ***z***_*u*_ ∼ *p*_*u*_(***z***_*u*_|***w***), where ***a*** stands for the domain index, ***w*** stands for the confounding, and we assume that ***z***_*d*_ ⊥ ***z***_*u*_|***w***.

The idea of counterfactual matching in the disentangled latent space is that, we assume that the effect of changing domains (perturbations) ***a*** on ***z***_*d*_ is rank-preserving, conditional on confoundings. That’s to say, perturbations might have heterogeneous effects on the latent factors of cells belonging to different cell types or cell states. However, after controlling for these confounding factors (cell type or cell state), the effect of the perturbation on the cells within that subgroup should be similar, meaning that the relative positions (distribution rank or quantile) of these cells before and after the perturbation should be preserved.

We resort to optimal transport (OT) for the implementation of quantile matching, i.e.,

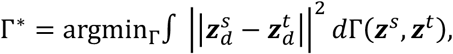

since there is an equivalence between OT with a strictly convex cost (like square Euclidean distance) and quantile matching in one-dimension [7, 8]. In multi-dimensional latent space, although inexact, we can view this as a trade-off in quantile matching across different dimensions, because the squared Euclidean distance can be seen as the summation of the squared Euclidean distances in each dimension.

When confounding ***w*** is not observed or not fully observed (e.g. only known a surrogate like cell type labels), ***z***_*u*_ can serve as a representation of confounding. It is hard to impose hard conditionals on ***z***_*u*_, i.e.,

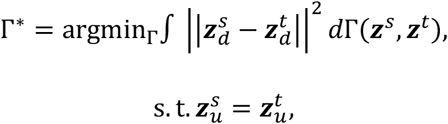

and we relax to a soft version,

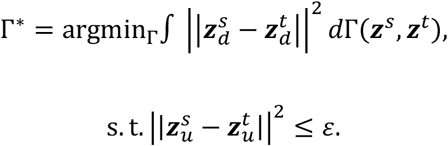

and this is equivalent to solving

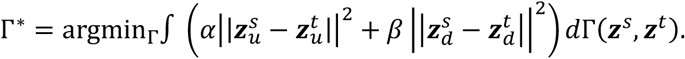

We can then generate counterfactual latent factors by barycentric projections on the full latent space

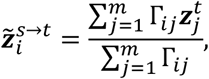

or on the ***z***_*d*_ space only

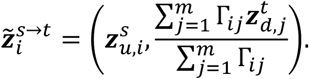

#### SN3 Toy examples to illustrate the idea of conditional optimal transport on disentangled latent space

We constructed synthetic datasets, ***X***_*s*_ (Source) and ***X***_*t*_ (Target), in a latent space. Each data point ***x*** = [***z***_*u*_, ***z***_*d*_] is composed of an **invariant component** (***z***_*u*_) and a **domain-specific component** (***z***_*d*_).

The ***z***_*u*_ component is designed to represent conserved cell identity (e.g., cell types), featuring two distinct, well-separated clusters, symmetrically centered around ± 2, ensuring that this fundamental structure is shared between ***X***_*s*_ and ***X***_*t*_. In contrast, the ***z***_*d*_ component models the domain shift or perturbation effect. While the source ***z***_*d*_ is centered at 0, the target ***z***_*d*_ is generated with cluster-specific shifts (± 2), resulting in a clear domain separation that is dependent on the underlying ***z***_*u*_ identity.

Specifically, the distributions are formulated as:

1. Invariant Factors (***z***_*u*_):

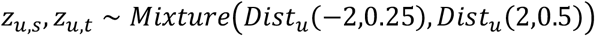
2. Domain-Specific Factors (***z***_*d*_):

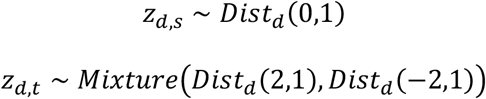

By allowing *Dist*_*u*_ and *Dist*_*d*_ to be selected from distributions such as Normal, Laplace, or Uniform, we can explore the robustness of the conditional optimal transport under various distributional differences, ensuring the generated data faithfully captures the need to align ***z***_*d*_ while strictly preserving the ***z***_*u*_ clustering structure.

We first conducted simulations in a setting where both ***z***_*d*_ and ***z***_*u*_ are 1-dimensional, and presented the scatter plots of the counterfactual generations ***x*** (Supplementary Fig.S1) and the marginal distributions of ***z***_*d*_ and ***z***_*u*_ (Supplementary Fig.S2). It can be seen that when no conditioning is performed (*α* = 0) or a very small *α* is set, significant cross-matching occurs between different groups. Although the marginals can be aligned, there is a clear issue of confounding interference. The issue becomes even more severe when performing barycentric mapping on ***z***_*d*_ only, as the lack of conditioning leads to a more severe distribution shift. Conversely, when ***z***_*u*_ is fully utilized for confounding matching without ***z***_*d*_ transfer (*β* = 0), we observe that while the marginals of ***z***_*u*_ can be aligned, the marginals of ***z***_*d*_ exhibit significant differences. The simulations conducted on a multi-dimensional latent space further corroborate this point (Supplementary Fig.S3). Therefore, using an appropriate level of conditioning is crucial.

**Supplementary Fig. S1.**
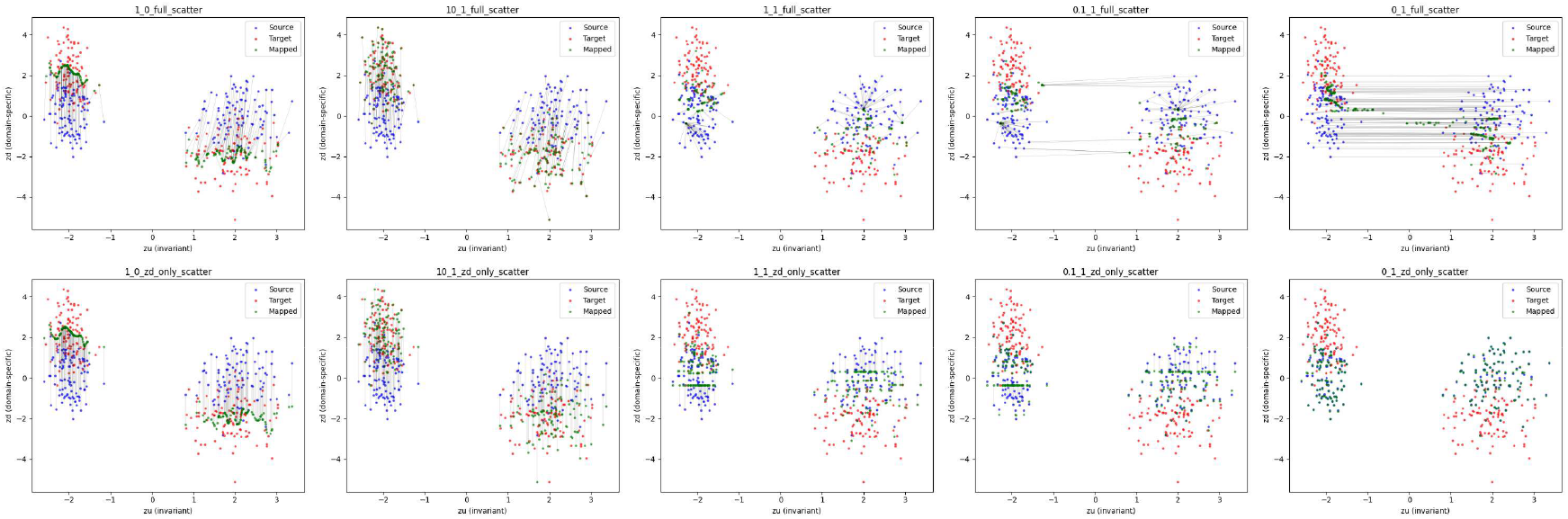
The scatter plots of the counterfactual generations in 1-dimensional latent-space toy example. The title of each subplot shows the configuration (alpha_beta_projection method) used.

**Supplementary Fig. S2.**
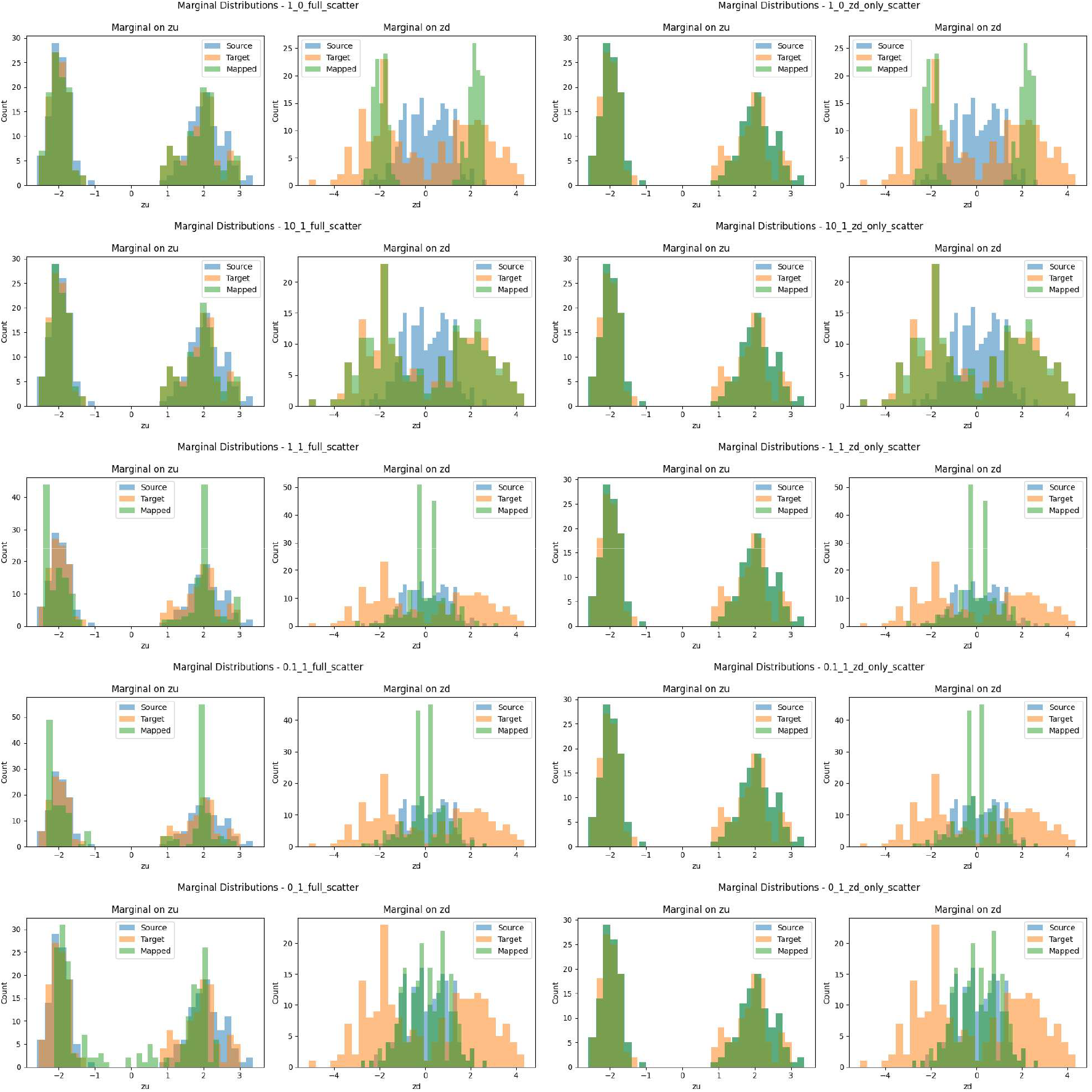
The marginal distributions of ***z***_*d*_ and ***z***_*u*_ in 1-dimensional latent-space toy example. The title of each subplot shows the configuration (alpha_beta_projection method) used.

**Supplementary Fig. S3.**
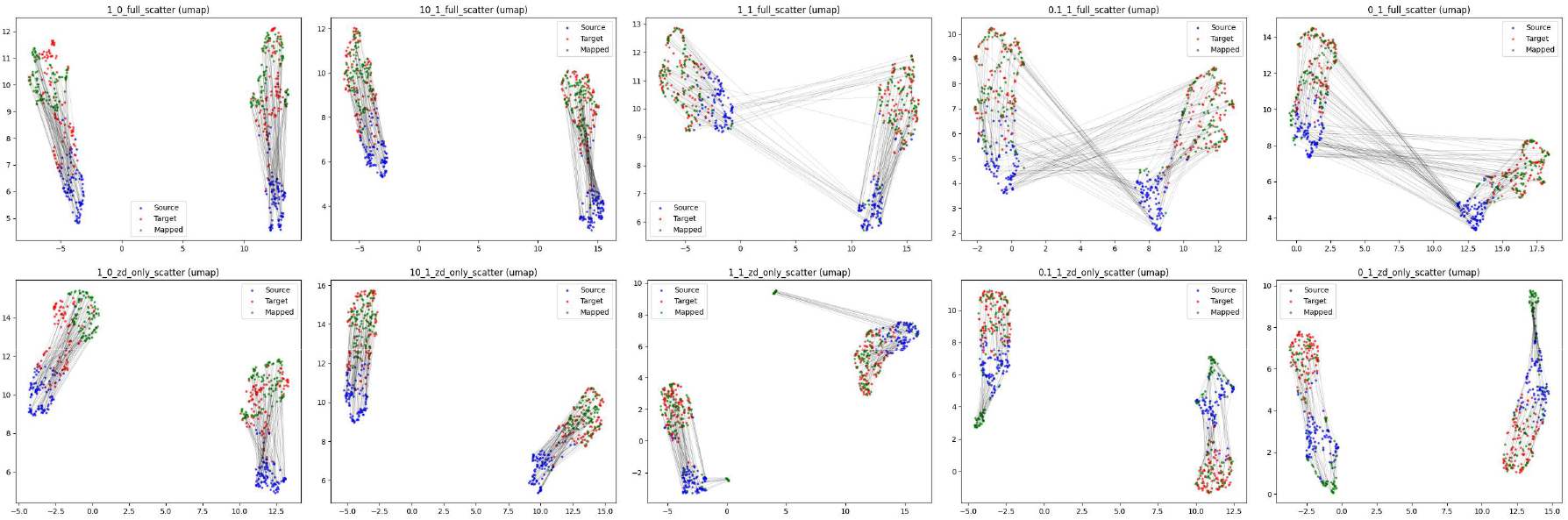
The scatter plots of the counterfactual generations in multi-dimensional latent-space toy example (*dim*(***z***_*d*_) = *dim*(***z***_*u*_) = 10 here). The title of each subplot shows the configuration (alpha_beta_projection method) used.

## SN4 Validating the control-to-treatment conditional rank-preserving correspondence

### Design

Following Methods Section 4.6, we simulated a small-size example data with 200 genes, in which each treated cell is generated as a constant, cell-type-specific additive shift *δ*_*c*_ of an individual control cell, with *z*_*u*_ unchanged. The same cell therefore exists in both states, giving a ground-truth one-to-one pairing (control cell *i* ↔ treated cell *i*) that destructive sequencing cannot provide. We form two populations: all cells in the control state, and the same cells in the treated state, and pass them to the conditional OT without any pairing information, then ask whether the recovered coupling re-pairs each control cell with its true counterpart.

### Coupling

For each cell type *c* (denote sets *S*_*c*_ for control, *T*_*c*_ for treated, where |*S*_*c*_| = |*T*_*c*_| = *n*_*c*_), we solve the conditional OT with cost

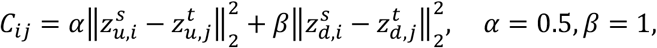

using the exact EMD solver, and row-normalise the plan so that ∑_*j*_ *Γ*_*ij*_ = 1 (barycentric weights). Matching is restricted within each cell type. In this paired setting the two populations have equal size, so the exact-EMD plan is a one-to-one permutation; in practice the control and treated populations differ in size (or an entropic/Sinkhorn solver is used), so the coupling is soft, i.e., each control cell is matched to a transport-weighted set of treated cells, and the matched score defined below is the barycentric average over that set.

### Response rank

The perturbation acts along the known shift direction 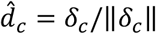. Each cell’s response score is the projection of its ground-truth *z*_*d*_ onto this axis, 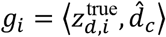, and its rank *r*_*i*_ is the percentile rank of *g*_*i*_ within its cell type. For a control cell *i*, the OT-matched score is the barycentric average 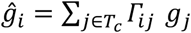, with matched rank 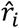 the percentile rank of 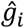

### Metrics (per cell type)

We use the Spearman rank correlation and top-1 index

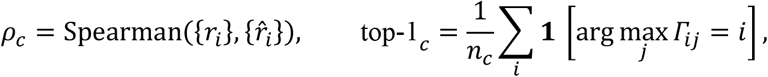

i.e. *ρ*_*c*_ measures whether a control cell is matched to a treated cell of the same response rank, and top-1_c_ the fraction of control cells whose highest-mass partner is their true counterpart *i*. Both are computed (i) on the ground-truth latent factors (we simulate *dim*(*z*_*d*_) = *dim*(*z*_*u*_) = 10) and (ii) on the factors inferred by the trained scDRP model (we set *dim*(*z*_*d*_) = *dim*(*z*_*u*_) = 20 with L0 penalty strength 0.01); in both cases the ranks 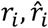 use the ground-truth *z*_*d*_ so that the reference ordering is exact and only the coupling *Γ* changes.

### Results

Shown in Table S1, on the ground-truth factors the within-cell-type coupling is the identity matrix (Fig. S4a) and rank preservation is perfect in all cell types (*ρ* = 1.00, top-1 = 1.00) — the unique optimum under a squared-Euclidean cost and a constant shift. On the model-inferred factors the correspondence is substantially preserved: mean per-cell-type *ρ* = 0.84 and mean top-1 partner recovery = 0.85, with the coupling remaining concentrated on the diagonal (Fig. S4b). The degree of preservation varies across cell types, reflecting how faithfully each cell type’s latent structure is recovered by the encoder. Together these results show that scDRP resolves the individual control-to-treatment correspondence, both in principle and through the learned representation.

**Supplementary Table S1.** Per-cell-type matching on simulated data. Spearman rank correlation (*ρ*) and top-1 partner recovery, on the ground-truth and model-inferred latent factors.

| cell type | n | $\rho$ (true) | top-1 (true) | $\rho$ (learned) | top-1 (learned) |
| --- | --- | --- | --- | --- | --- |
| 0 | 230 | 1.00 | 1.00 | 0.78 | 0.88 |
| 1 | 264 | 1.00 | 1.00 | 0.63 | 0.60 |
| 2 | 200 | 1.00 | 1.00 | 0.88 | 0.84 |
| 3 | 244 | 1.00 | 1.00 | 0.97 | 0.99 |
| 4 | 312 | 1.00 | 1.00 | 0.96 | 0.95 |
| **mean** | — | **1.00** | **1.00** | **0.84** | **0.85** |

**Supplementary Fig. S4.**
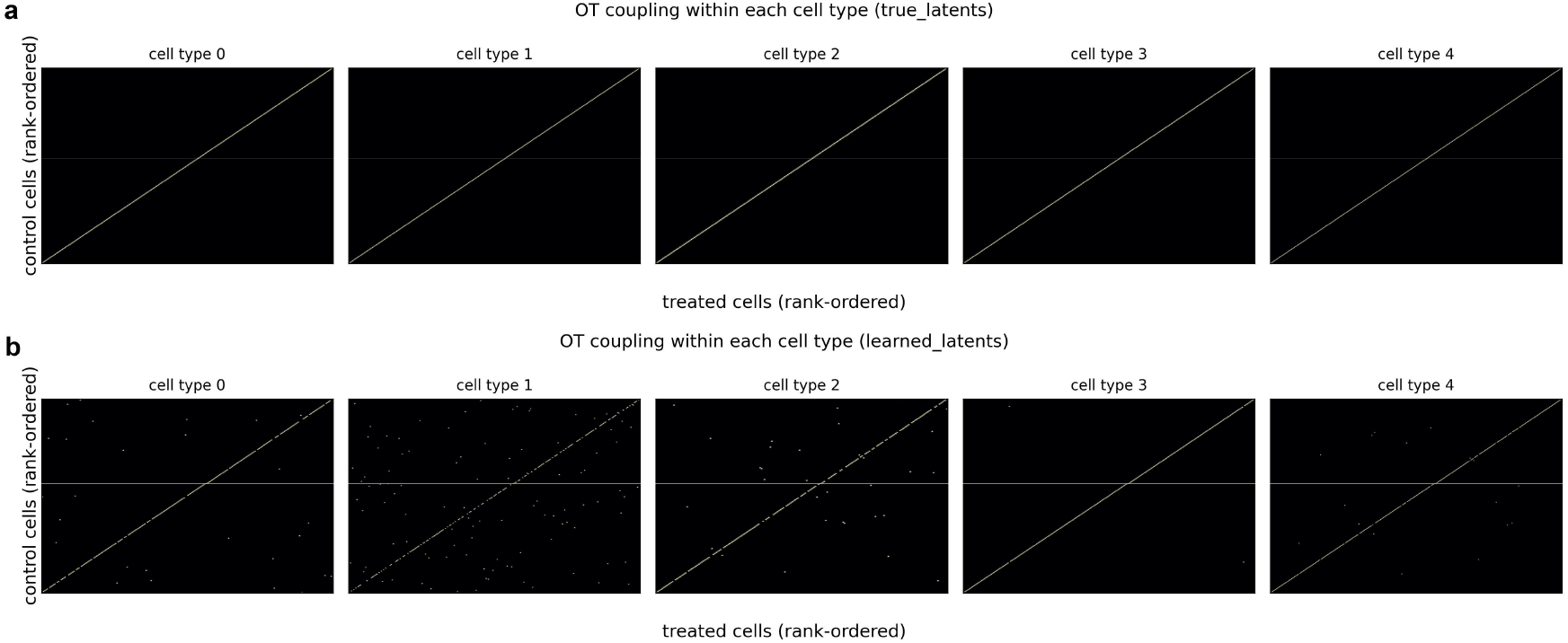
Within-cell-type OT coupling Γ (control rows vs treated columns, both ordered by response rank; one panel per cell type) on (**a**) the true latent factors in data generating process and (**b**) the model-inferred latent factors. A bright entry at position (*m, m*) indicates that the control cell of rank *m* is matched to the treated cell of the same rank; the coupling is the identity in (a) and remains concentrated on the diagonal in (b).

## SN5 Visualization on learned latent representations from real dataset

To evaluate the efficacy of the learned disentangled latent representations, we performed UMAP visualizations on the Kang dataset for the perturbation-induced latent space (*z*_*d*_), the perturbation-invariant latent space (*z*_*u*_), and the full latent representation (*z*), across two experimental settings: with and without the provision of cell-type labels.

It can be observed that the perturbation-invariant space *z*_*u*_ effectively isolates perturbation-invariant, intrinsic cellular characteristics. Specifically, cells from the treatment and control groups are integrated together, whereas different cell types are distinctly clustered (Fig. S5a). Notably, this precise clustering of cell types is maintained even when cell-type labels are not provided in the training process (Fig. S6a). In contrast, the perturbation-induced space *z*_*d*_ segregates the treatment group from the control group, while simultaneously distinguishing between cell types due to their heterogeneous cell-type specific perturbation responses (Fig. S5b, S6b). When visualizing the full latent space, the UMAP projections clearly indicate that both cell types and experimental conditions are well-resolved, characterized by two separate clusters (treatment and control) for each individual cell type (Fig. S5c, S6c).

Consequently, these results validate that our disentangled representation learning method accurately captures and decouples the latent features underlying single-cell perturbation data, which are subsequently utilized in conditional OT to achieve precise causal effect identification and perturbation response prediction.

**Fig. S5.**
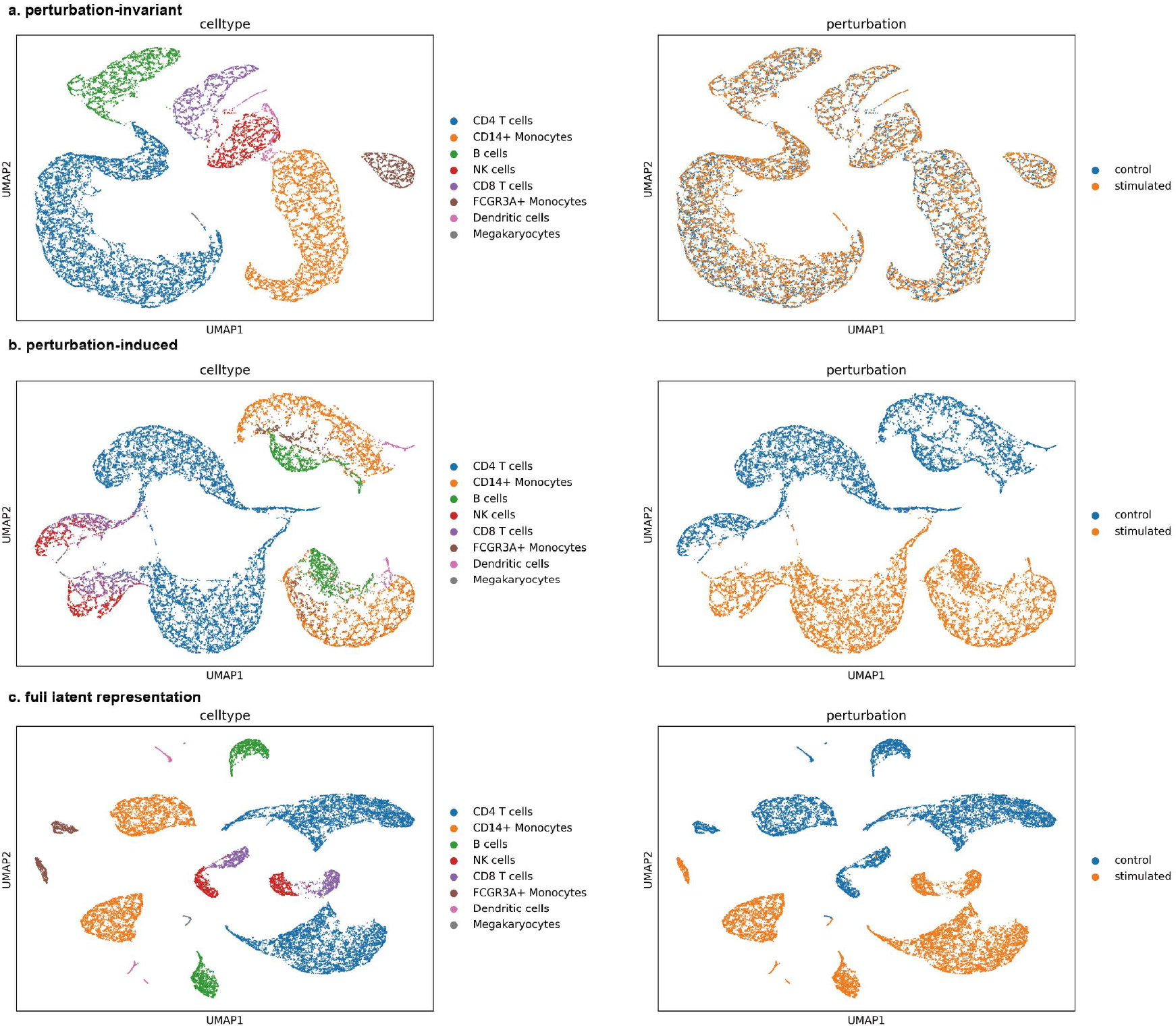
UMAP visualization of learned latent perturbation-invariant (*z*_*u*_, a), perturbation-induced (*z*_*d*_, b) and full latent representation (*z*, c) for Kang dataset, where the original cell-type labels are explicitly provided in the training process.

**Fig. S6.**
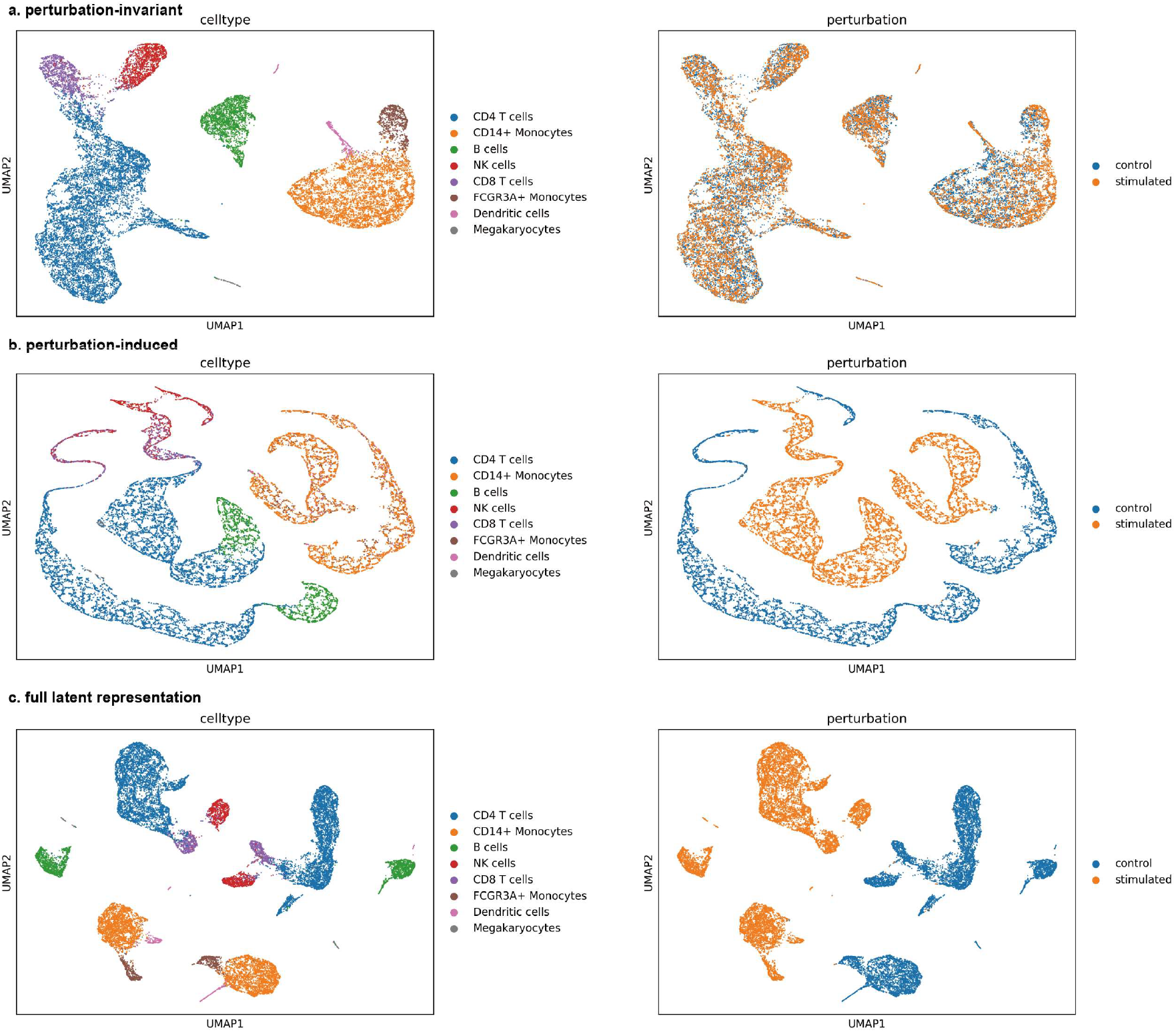
UMAP visualization of learned latent perturbation-invariant (*z*_*u*_, a), perturbation-induced (*z*_*d*_, b) and full latent representation (*z*, c) for Kang dataset, where the original cell-type labels are not provided in the training process. We can see the representation learning process can still recover the accurate cell identity structures.

---

**Supplementary Figure S7.**
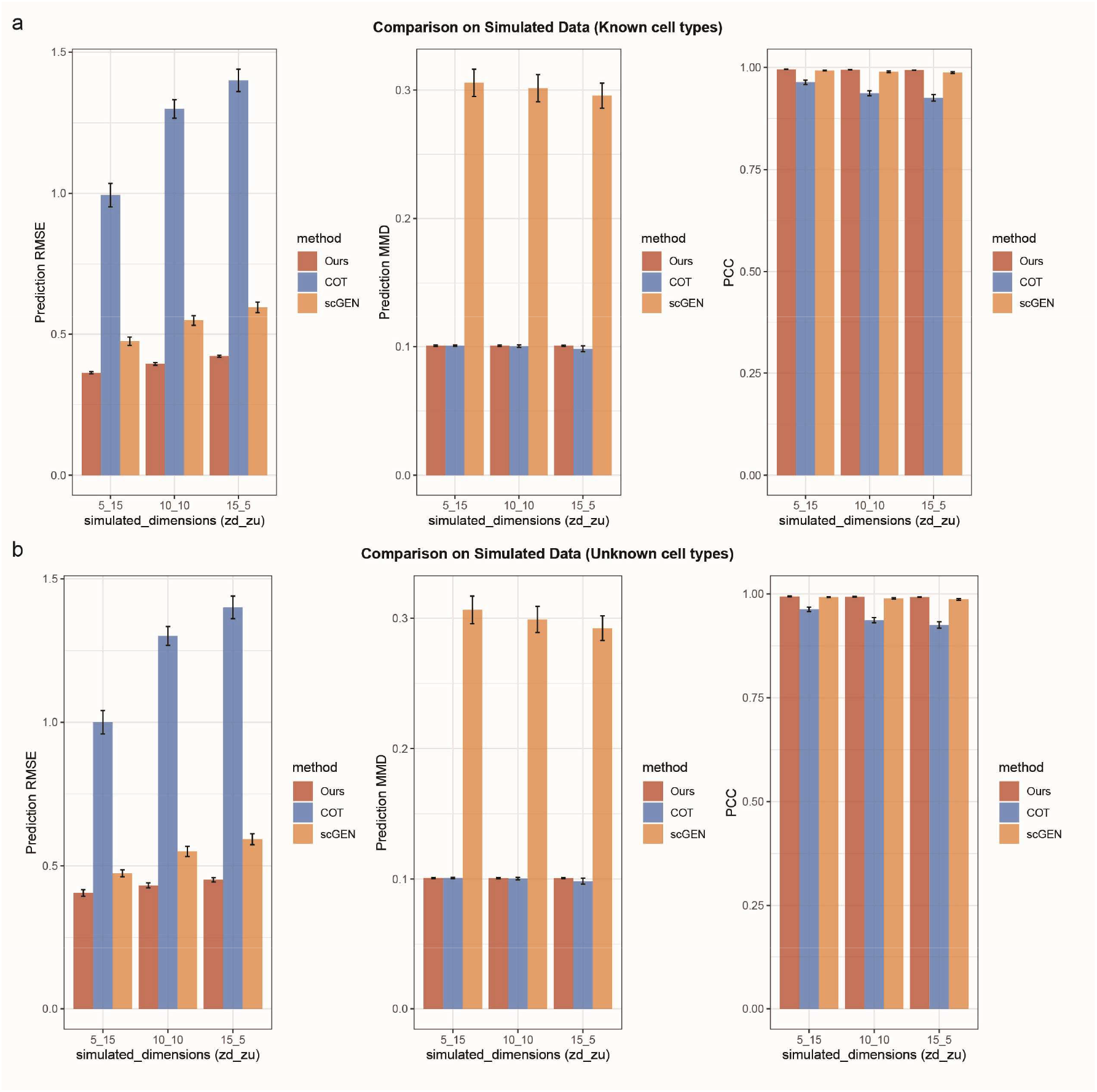
Performance on counterfactual predictions with simulated data under known cell-type labels (a) and unknown cell-type labels (b) settings. We evaluated the RMSE and Pearson correlation coefficient (PCC) of the mean expression of each gene between observed treatment group and counterfactual predictions from control group, as well as the maximum mean discrepancy (MMD) between the observed and predicted distributions. All metrics are averaged across all perturbations and cell types in each simulation. The x-axis shows the dimension settings for *z*_*d*_ and *z*_*u*_ in the simulation, and the different colors correspond to different methods. We repeat each simulated setting 20 times and report the mean±1.96SE.

**Supplementary Figure S8.**
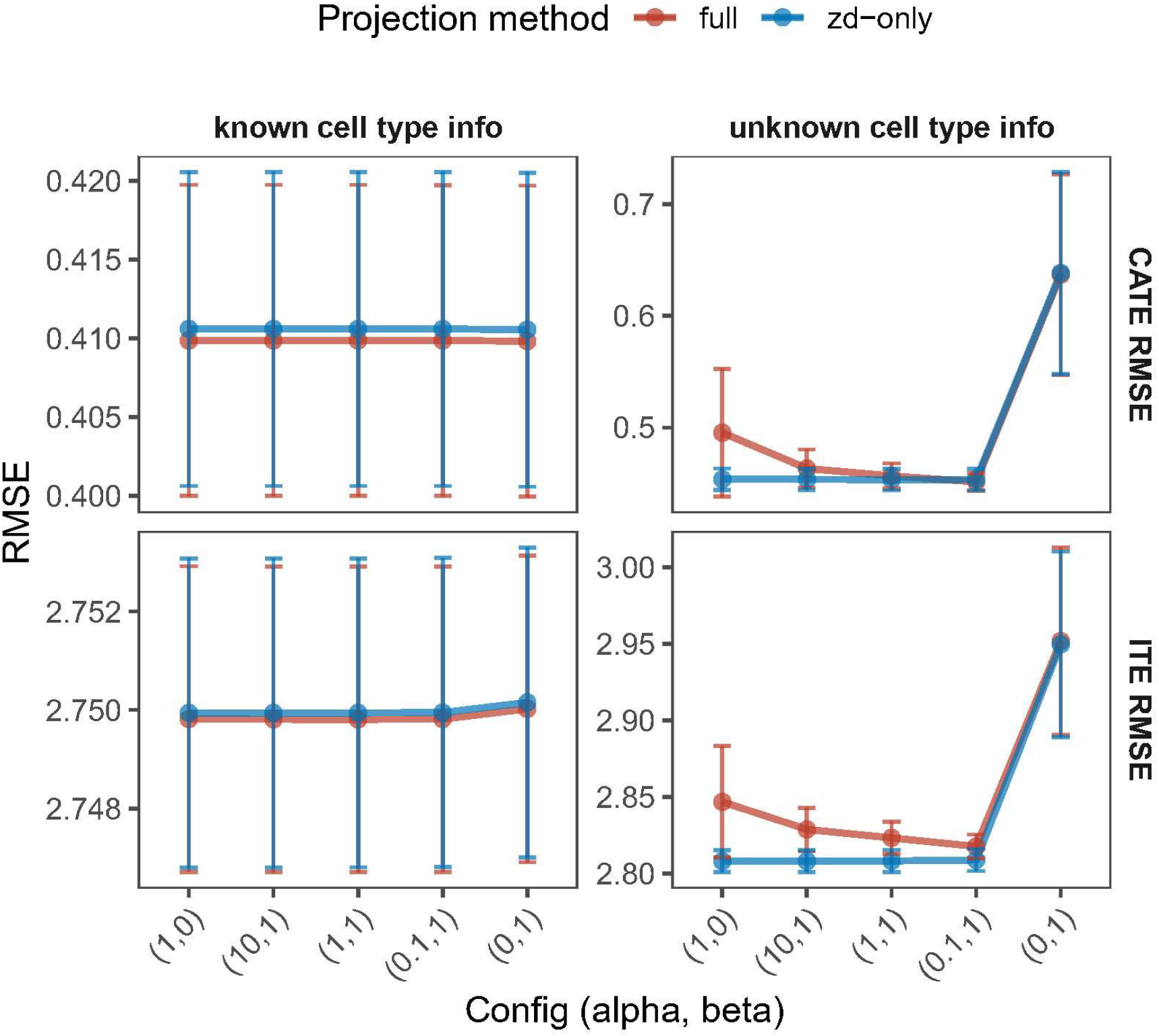
Sensitivity analysis on hyperparameters of conditional OT on simulated data. In each hyperparameter configuration, we repeat each simulated setting 20 times and report the mean RMSE±1.96SE.

**Supplementary Figure S9.**
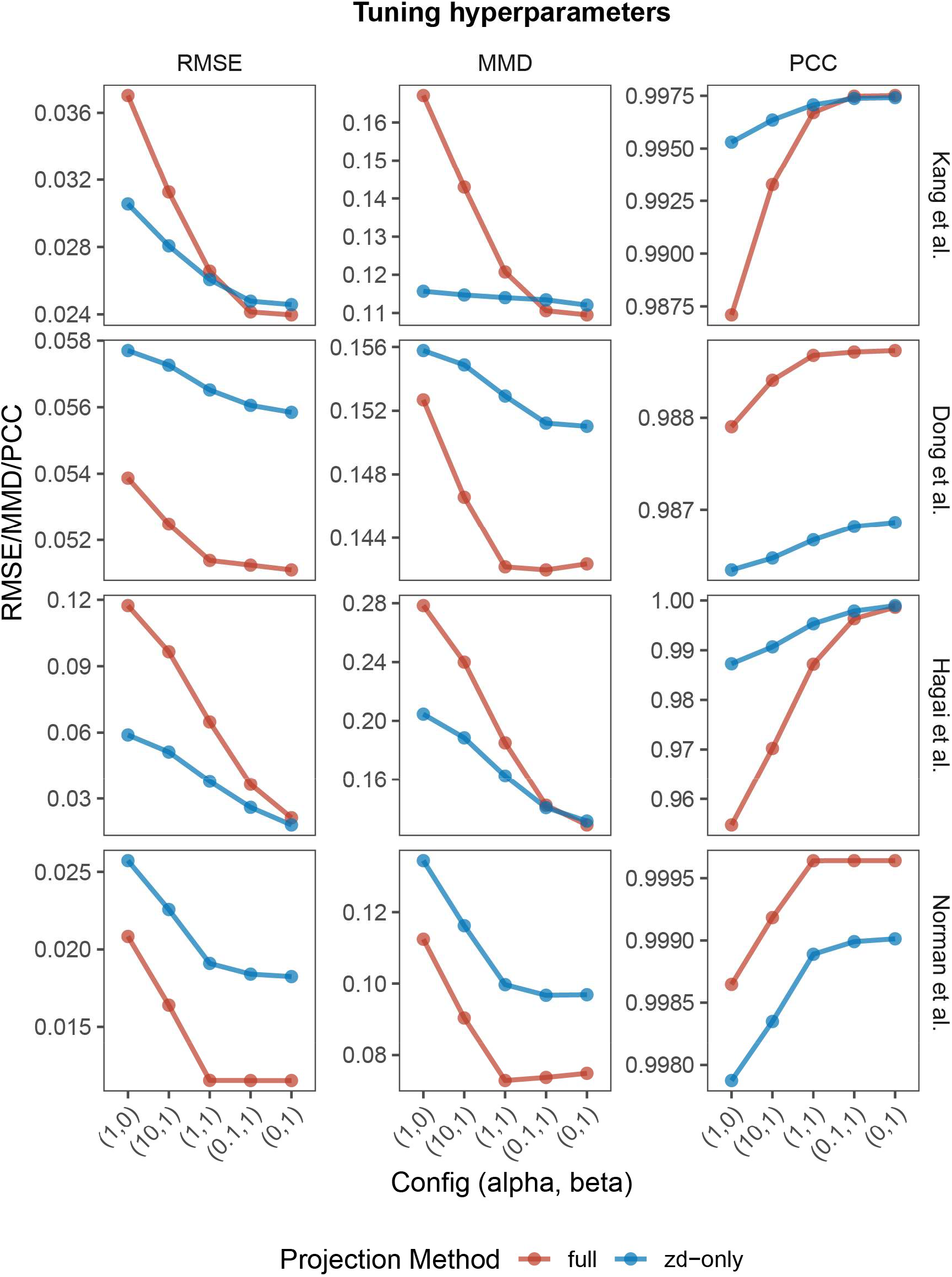
Sensitivity analysis on hyperparameters of conditional OT on real data (known cell-type label).

**Supplementary Figure S10.**
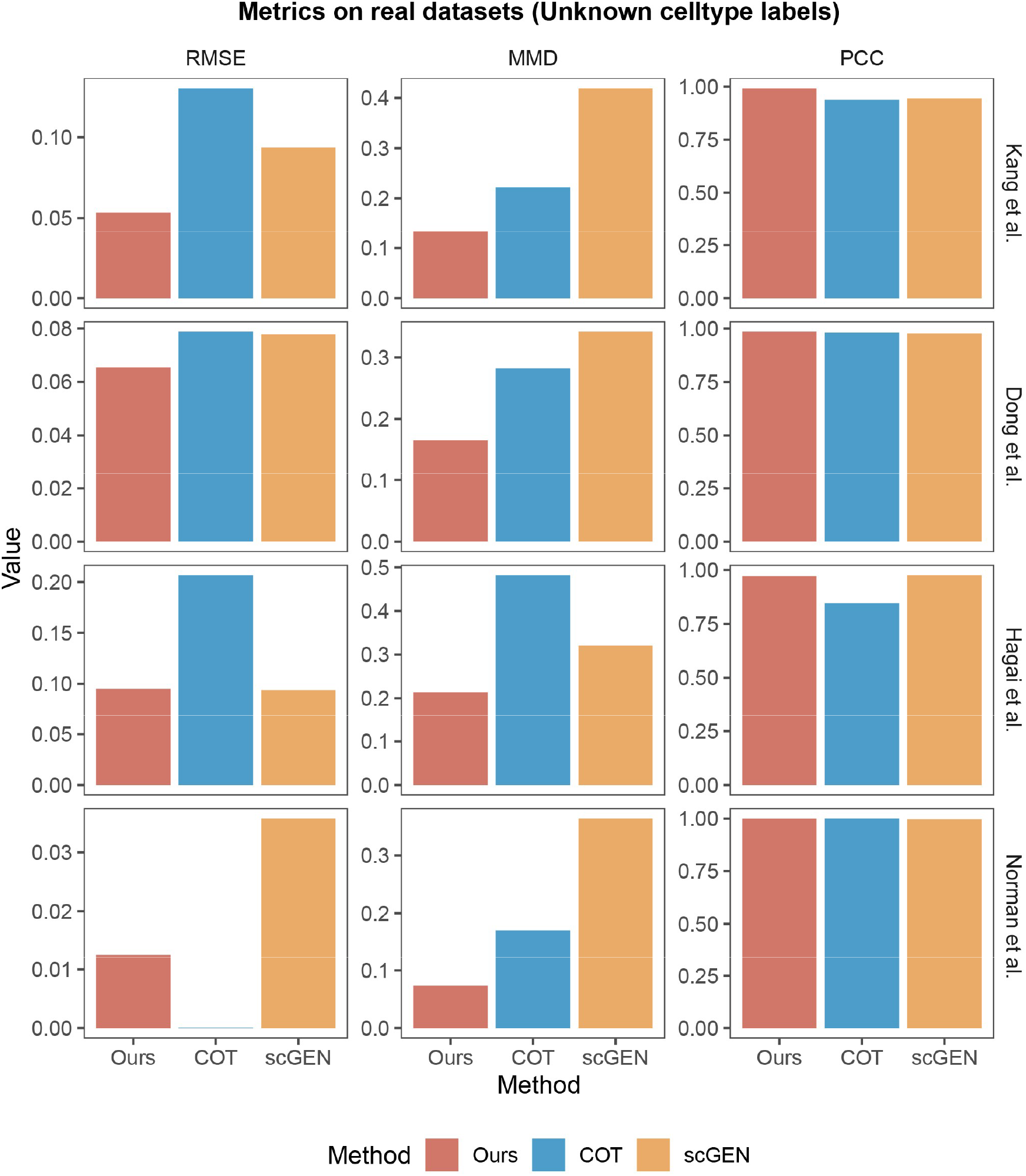
Performance on counterfactual predictions on real single-cell data without cell-type label information provided in training. The evaluation metrics includes RMSE and PCC of the mean expression of each gene between observed treatment group and counterfactual predictions from control group (we compare the mean since the true counterfactual outcome of control cells are not accessible in real data), as well as MMD between the observed and predicted distributions. All metrics are averaged across all perturbations and cell types.

**Supplementary Figure S11.**
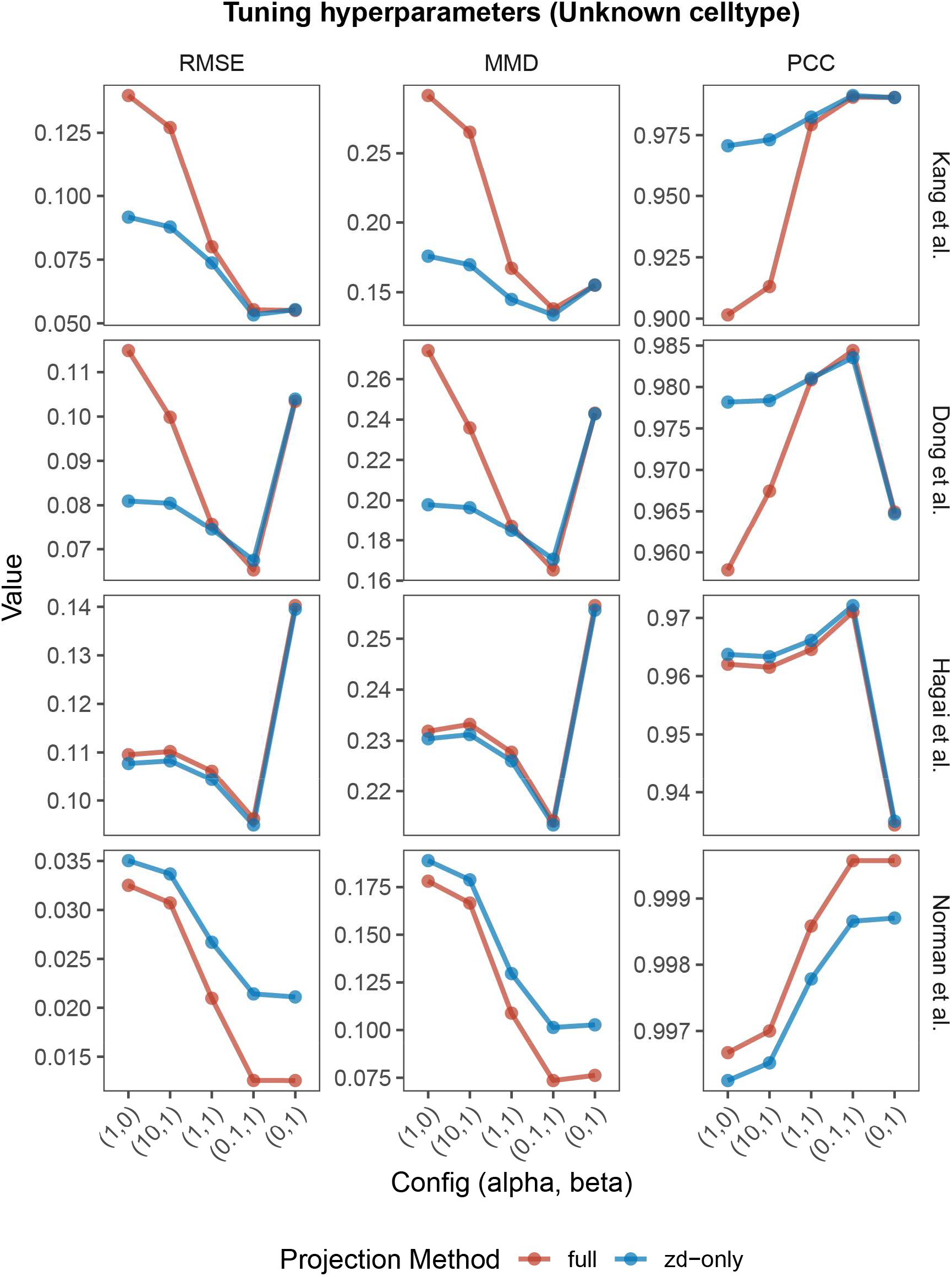
Sensitivity analysis on hyperparameters of conditional OT on real data under the unknown cell-type labels setting.

**Supplementary Figure S12.**
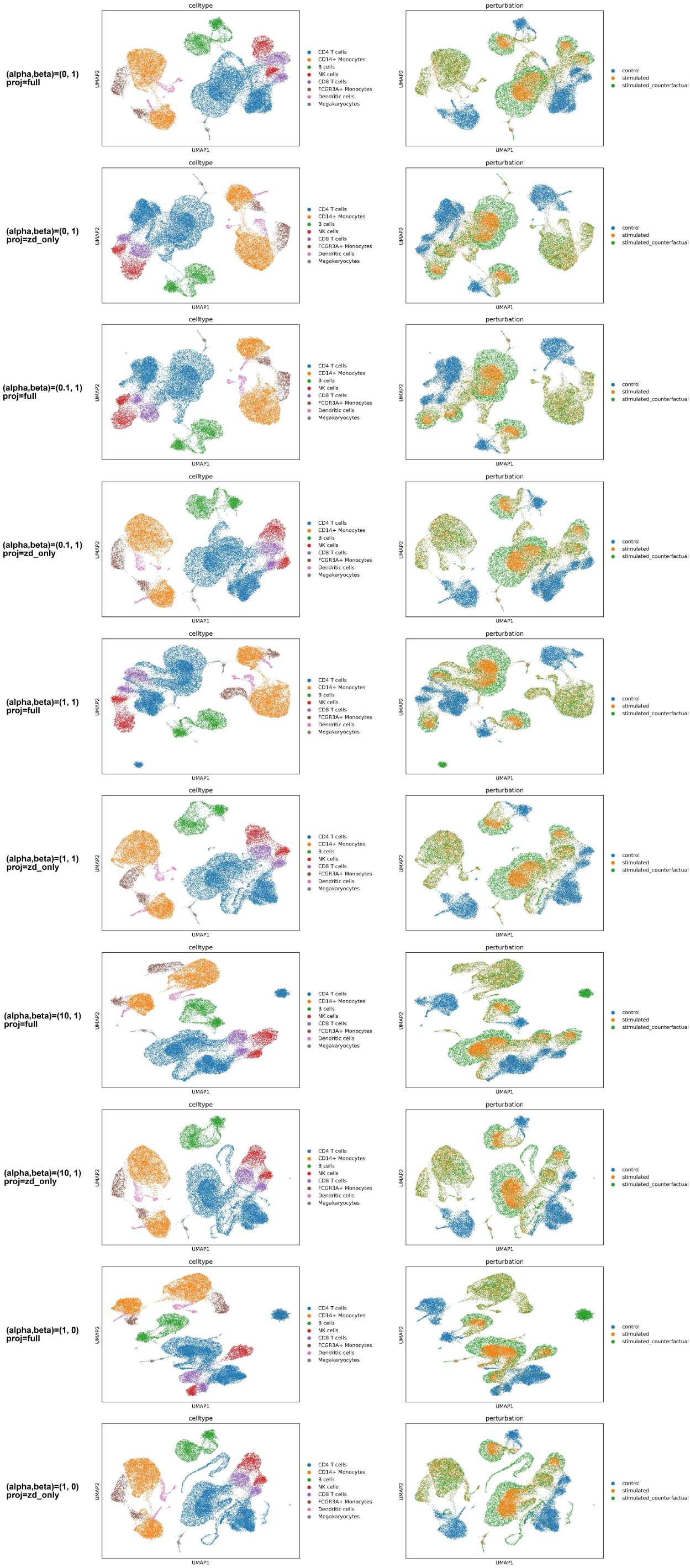
UMAP visualization of counterfactual predictions on Kang dataset under different settings of alpha, beta, and barycentric projection methods (known cell types).

**Supplementary Figure S13.**
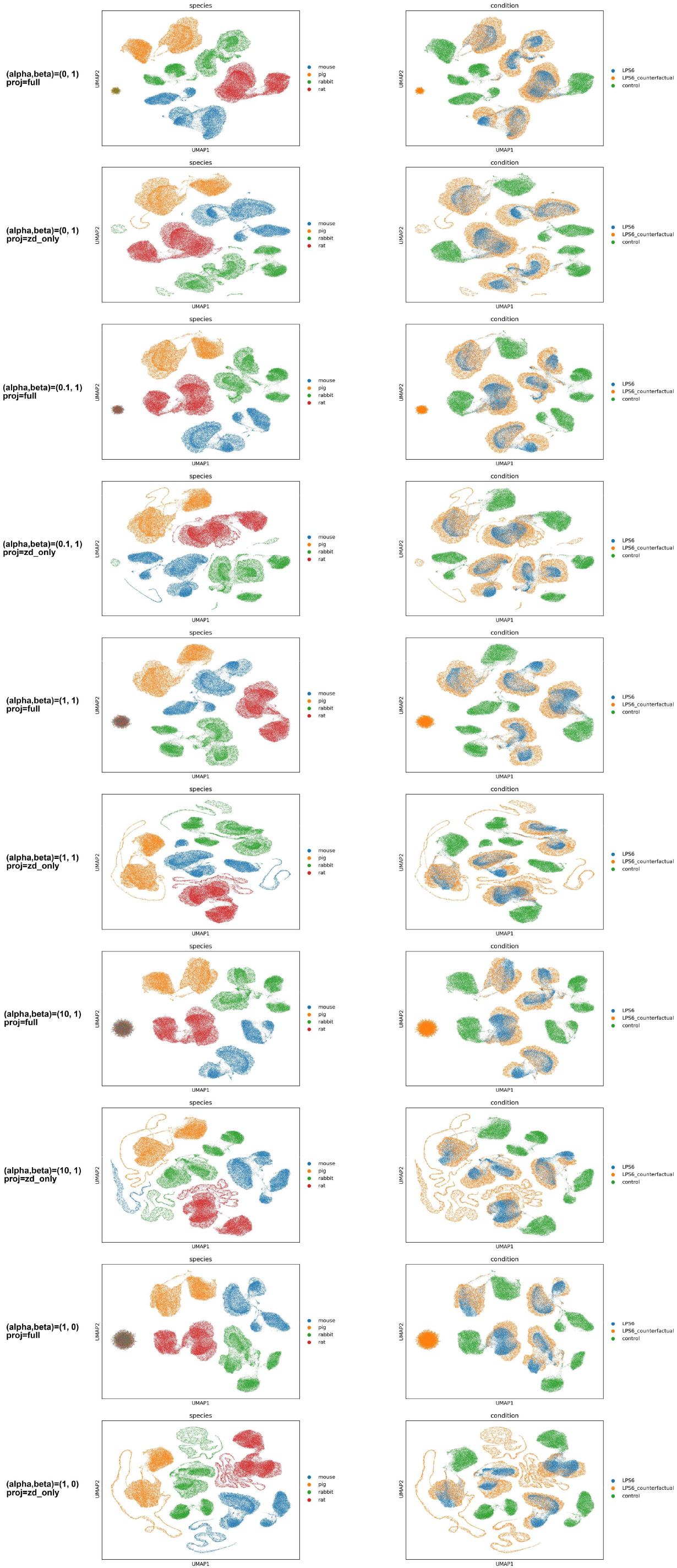
UMAP visualization of counterfactual predictions on Hagai dataset under different settings of alpha, beta, and barycentric projection methods (known cell types).

**Supplementary Figure S14.**
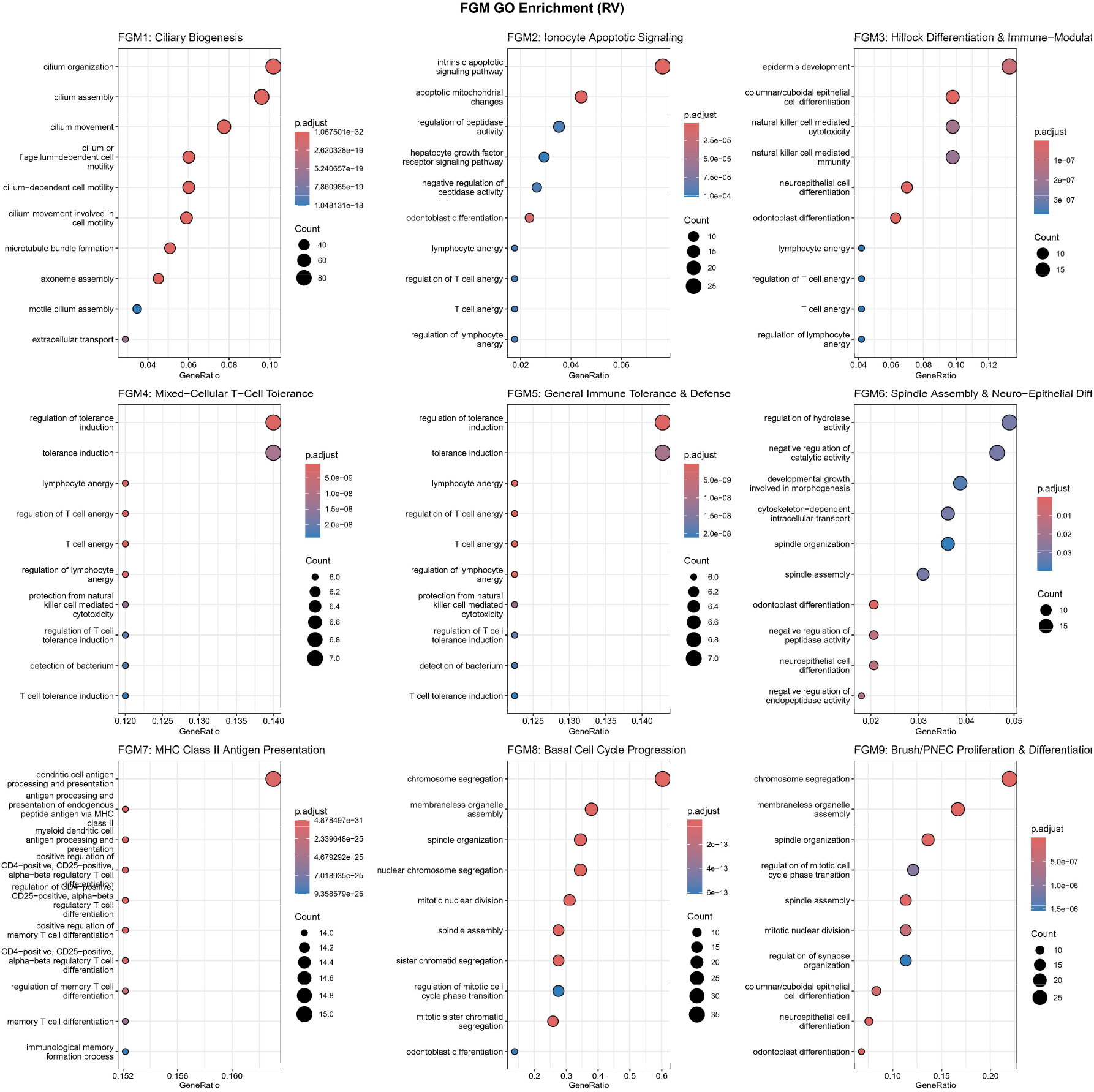
Details on GO enrichment of FGM identified in primary human bronchial organoids under rhinovirus (RV) exposure.

**Supplementary Figure S15.**
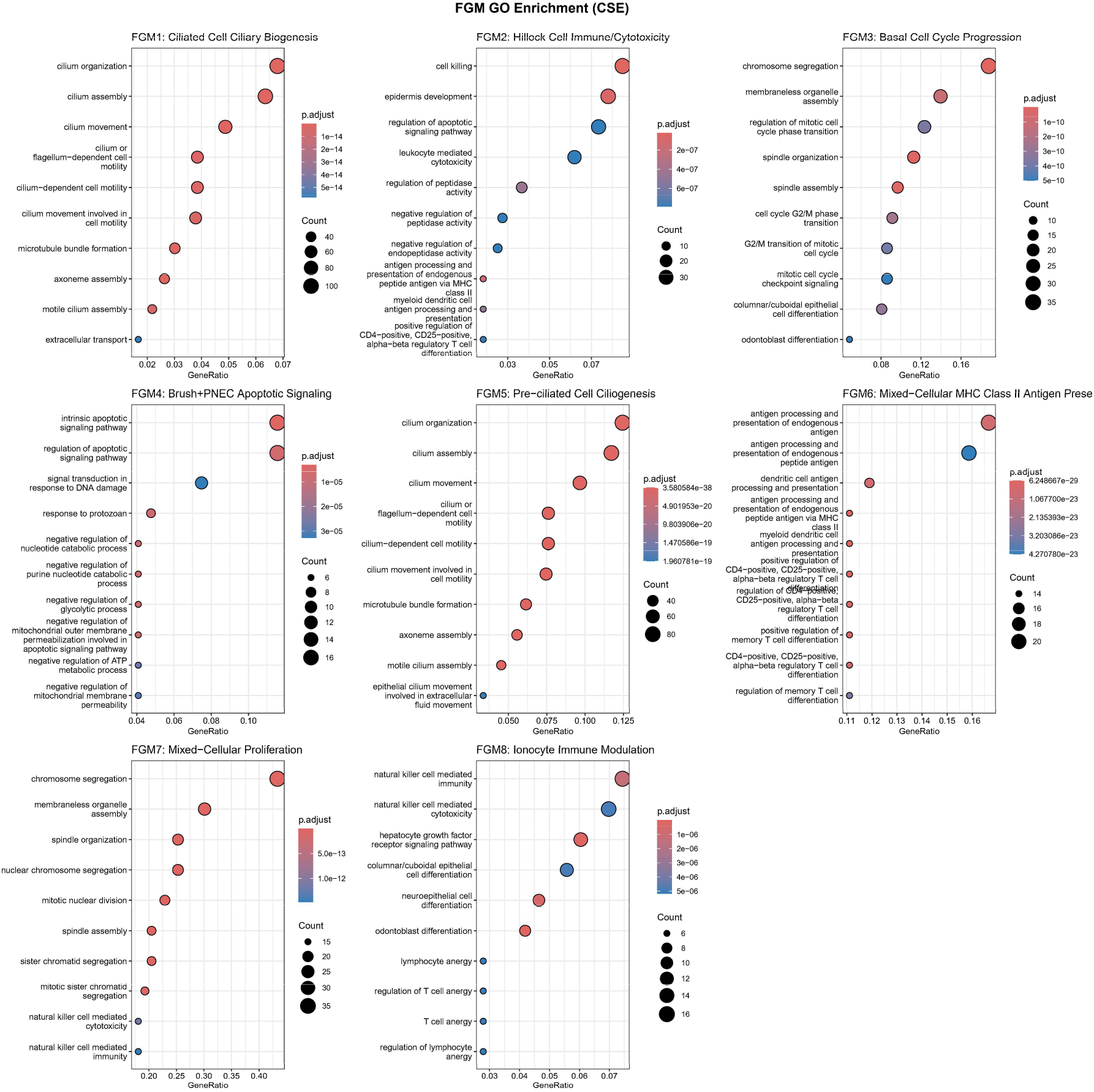
Details on GO enrichment of FGM identified in primary human bronchial organoids under cigarette-smoke extract (CSE) exposure.

**Supplementary Figure S16.**
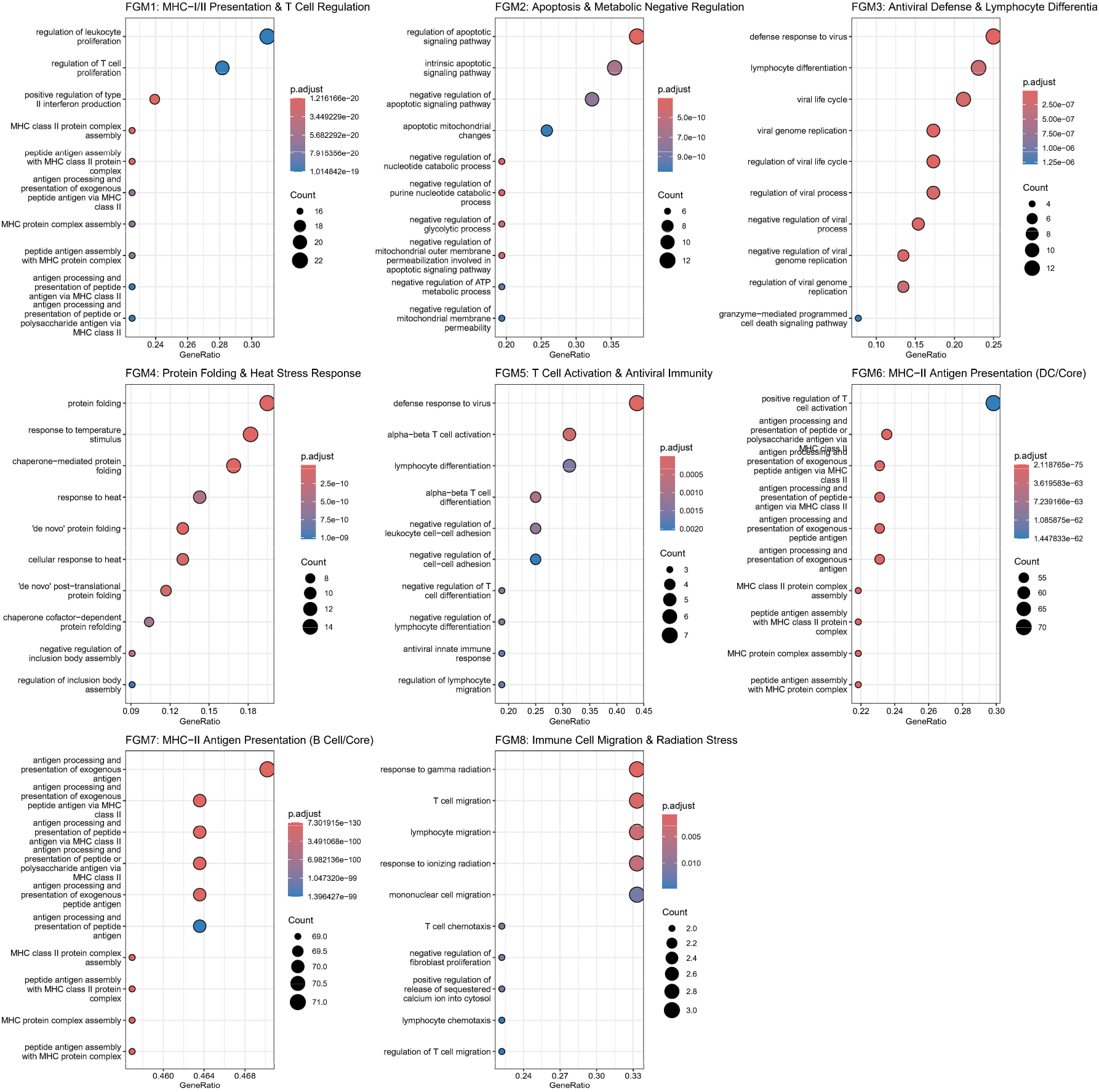
Details on GO enrichment of FGM identified from Kang dataset.

**Supplementary Figure S17.**
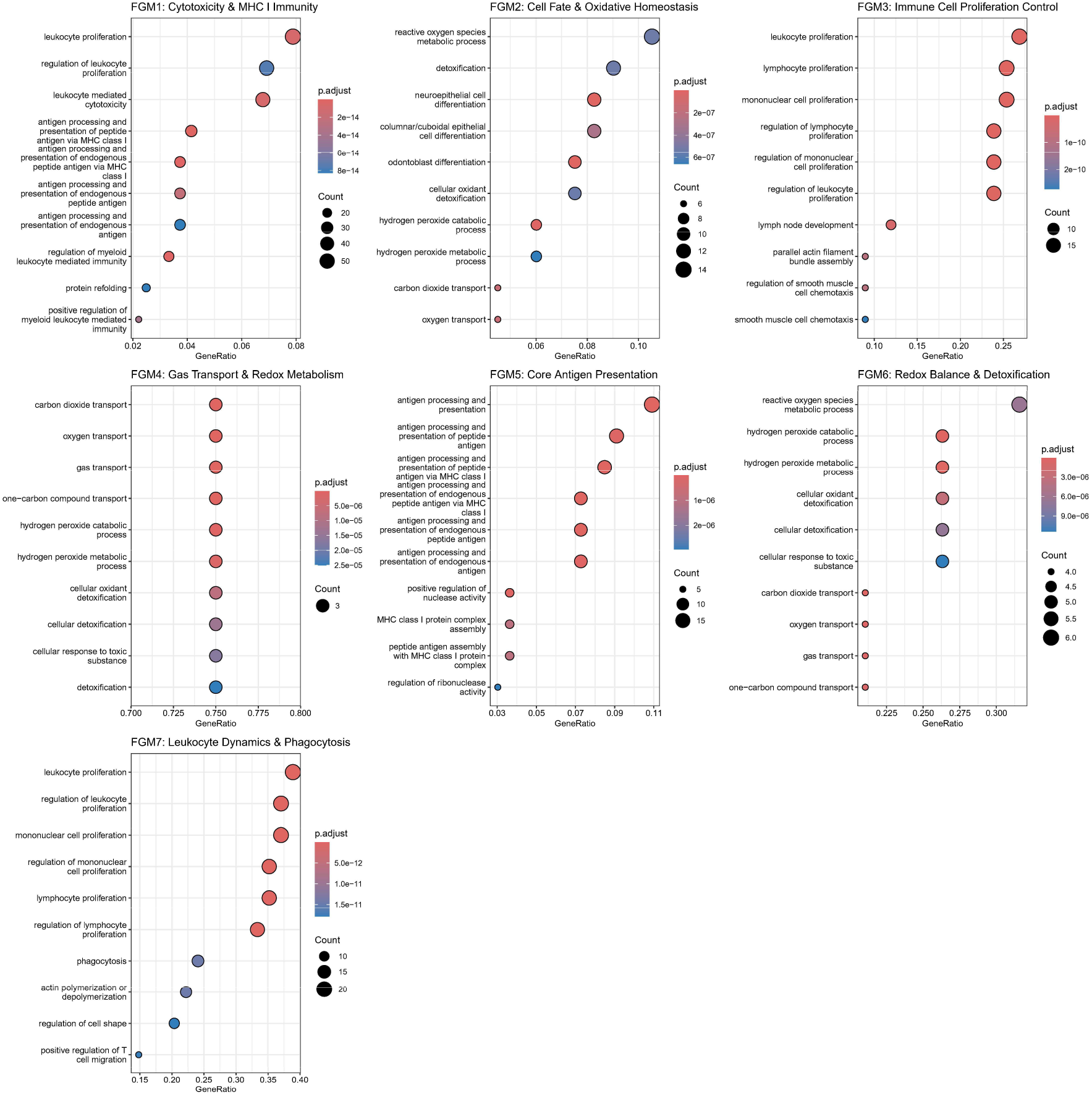
Details on GO enrichment of FGM identified from Norman CRISPR-a Perturb-seq dataset.

## Supplementary Note SN6 Nucleosome-related gene effect analysis with CRISPR-a Perturb-Seq

We investigated how CRISPR-a perturbations relate to nucleosome-related and non-nucleosome-related genes using the estimated ITE matrix on the Norman data. We used CpG island (CGI) promoter status as an imperfect proxy for promoter nucleosome architecture: non-CGI promoters tend to be assembled into stable, positioned nucleosomes and to require stimulus-dependent chromatin remodeling for activation, whereas CGI promoters tend to be nucleosome-depleted and constitutively accessible [9,10]. This proxy is grounded in the sequence determinants of nucleosome organization: CpG and GC content correlate with nucleosome depletion at mammalian promoters both *in vivo* and *in vitro* [11], and interpretable sequence-based models trained on MNase-seq data identify CpG-containing patterns among the dominant determinants of nucleosome organization: the CCG/CGG trinucleotide is depleted on nucleosomal relative to inter-nucleosomal DNA, and its periodicity is the top-ranked feature for discriminating nucleosomal from inter-nucleosomal regions [12]. CGI status can therefore be viewed as a coarse, promoter-level summary of the same sequence signal that dictates nucleosome organization at nucleotide resolution.

Each gene was classified by overlapping its promoter region (TSS±1 kb) with annotated CpG islands, yielding 942 non-CGI and 1,045 CGI genes out of the 3,000 highly variable genes. For each gene we computed its mean absolute ITE (effect magnitude) per perturbation from the inferred ITE matrix, and compared the two promoter classes both unadjusted (Mann– Whitney U test) and controlling for baseline expression (using expression as a covariate, as well as a more robust expression-decile-stratified test), given that both effect magnitude and CGI status correlate with expression level.

The result is an expression-dependent pattern. Without adjustment, CGI genes appear to have larger effects (Cliff’s δ ≈ ™0.2), but this is attributable to their higher baseline expression. After controlling for expression, the relationship reversed and remained consistent across all eight perturbations: non-CGI promoters show larger perturbation-induced transcriptional responses than CGI promoters, with a positive adjusted coefficient in all eight perturbations (expression-decile-stratified p < 0.01 for all eight; the expression-covariate model agrees in direction for all eight and reaches p < 0.05 for seven of eight) (Fig. S18). This is consistent with previous reports that non-CGI promoters generally display higher nucleosome occupancy and stronger nucleosome positioning, requiring stimulus-dependent chromatin remodeling for activation, whereas CGI promoters tend to adopt nucleosome-depleted and more permissive chromatin architectures [9, 10].

We acknowledge several key limitations of the current analysis. First, the effects, while statistically robust, are modest in magnitude. Second, CGI status is only an imperfect proxy for nucleosome occupancy: low nucleosome occupancy is a general property of active promoters and non-CGI promoters are heterogeneous. This imperfect proxy may reduce our ability to fully resolve the relationship between promoter chromatin architecture and perturbation responsiveness. Third, sequence-governed nucleosome positioning is statistical in nature and can be overridden *in vivo* by chromatin remodelers, as reflected in the reduced accuracy of sequence-based predictors in transcriptionally active relative to resting cells [12]. Analyses of perturbation effects on directly measured chromatin-level features, such as scATAC-seq data, remain open for future studies.

**Supplementary Figure S18.**
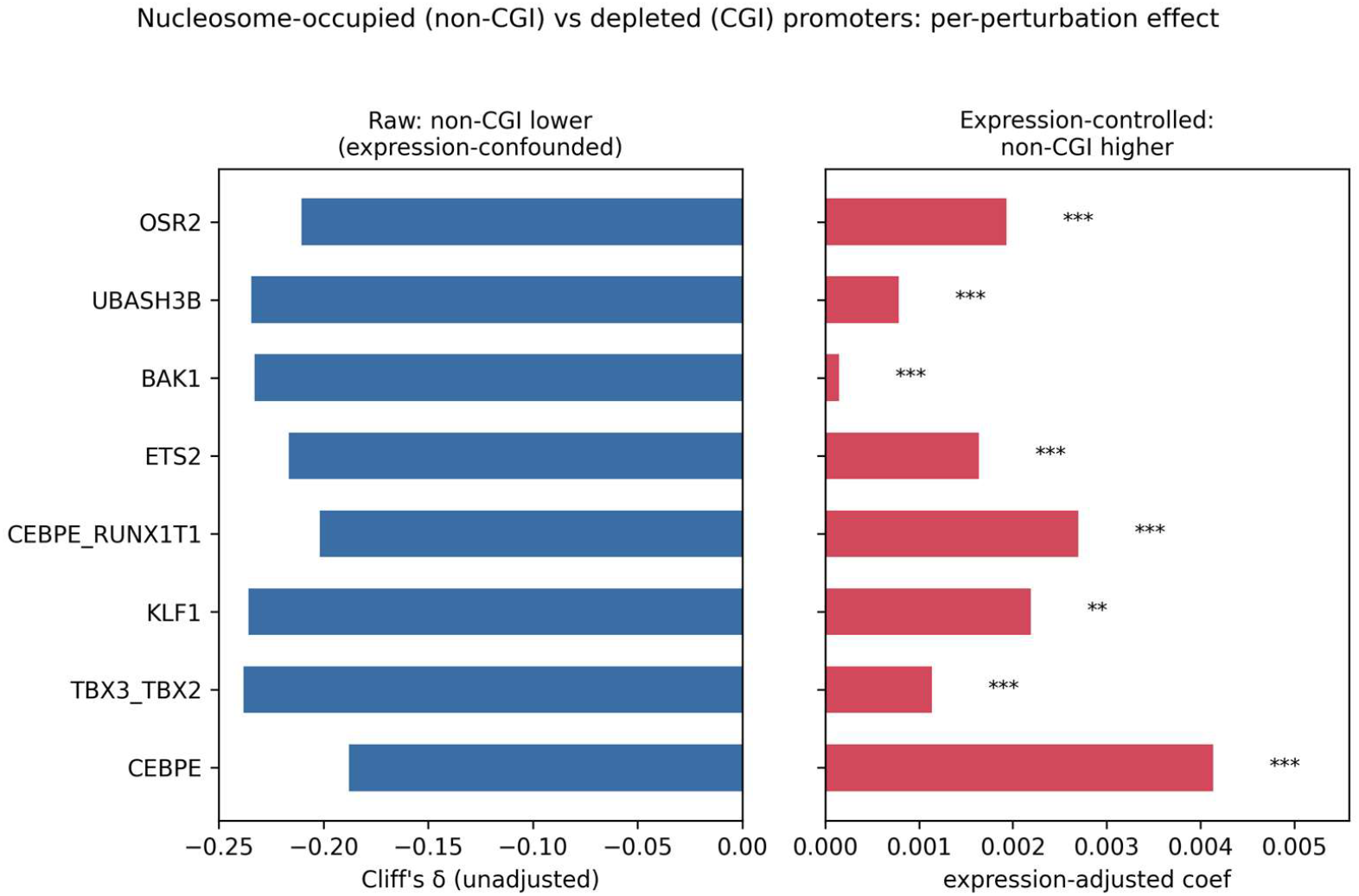
Nucleosome-occupied (non-CGI) promoter genes show larger treatment effects than nucleosome-depleted (CGI) genes after controlling for expression. Per-gene effect magnitude (mean |ITE|) was compared between genes with non-CGI (nucleosome-occupied, *n* = 942) and CGI (nucleosome-depleted, *n* = 1,045) promoters across the eight K562 CRISPR-activation perturbations. Left: unadjusted (Cliff’s δ); *δ* ≈ ™0.2 indicates non-CGI genes appear less responsive, reflecting their lower baseline expression. Right: after controlling for expression, the coefficient is positive in every perturbation: non-CGI genes are more responsive at matched expression (*** p < 0.001, ** p < 0.01, expression-decile-stratified test). CpG-island status is a proxy for promoter nucleosome architecture.

---

**Supplementary Table S2.**
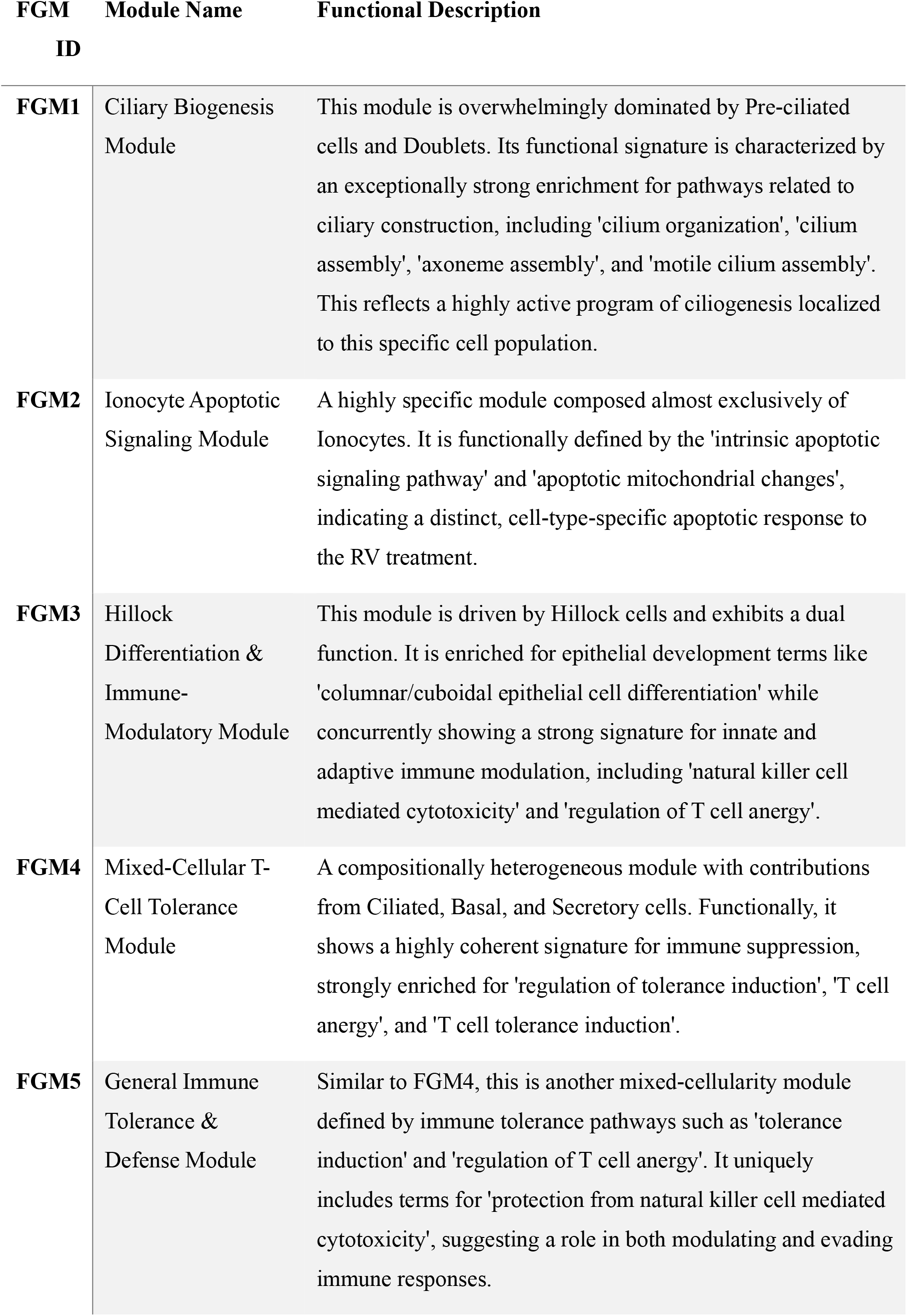
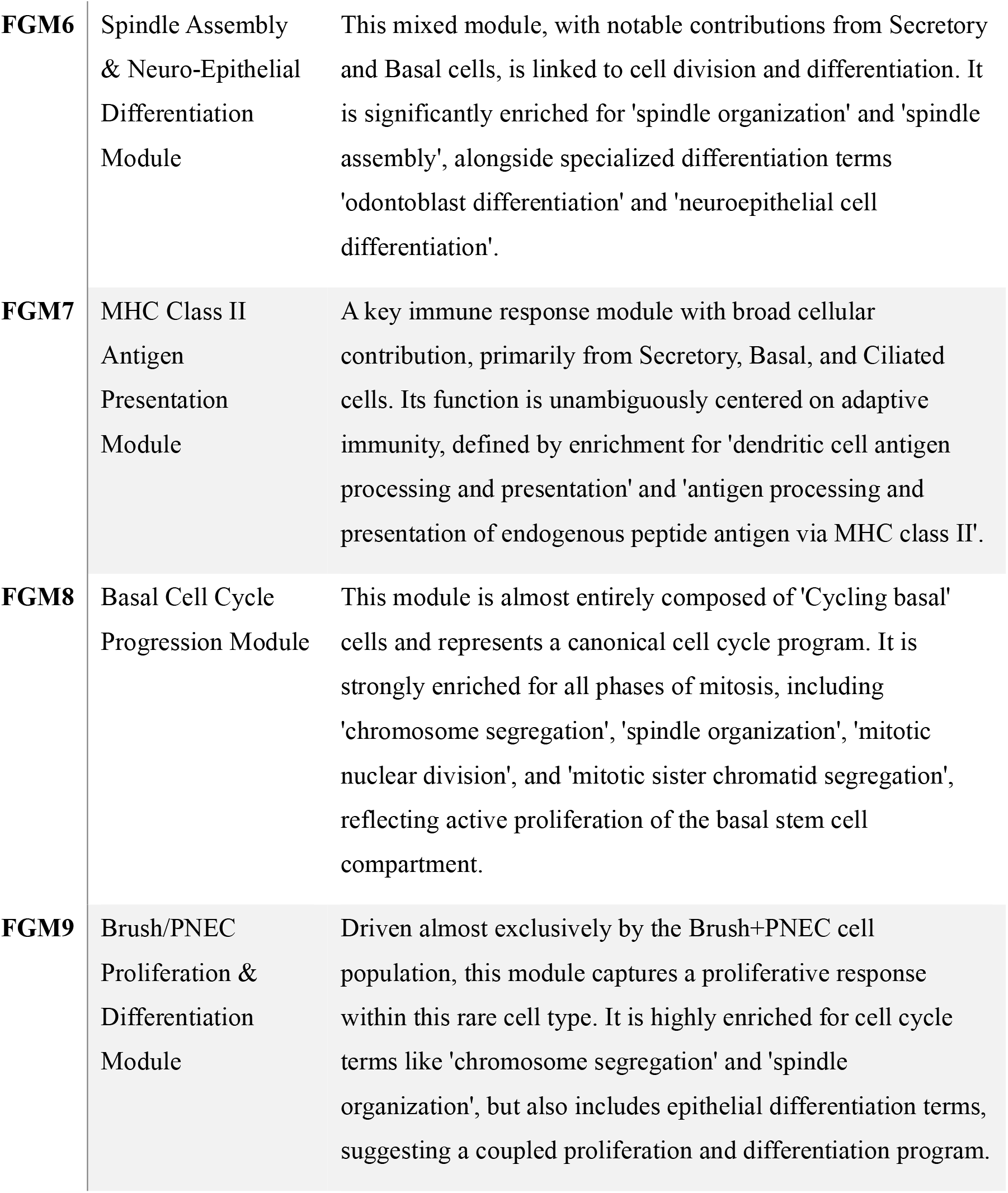
Details on the FGM identified in primary human bronchial organoids under rhinovirus (RV) exposure.

**Supplementary Table S3.**
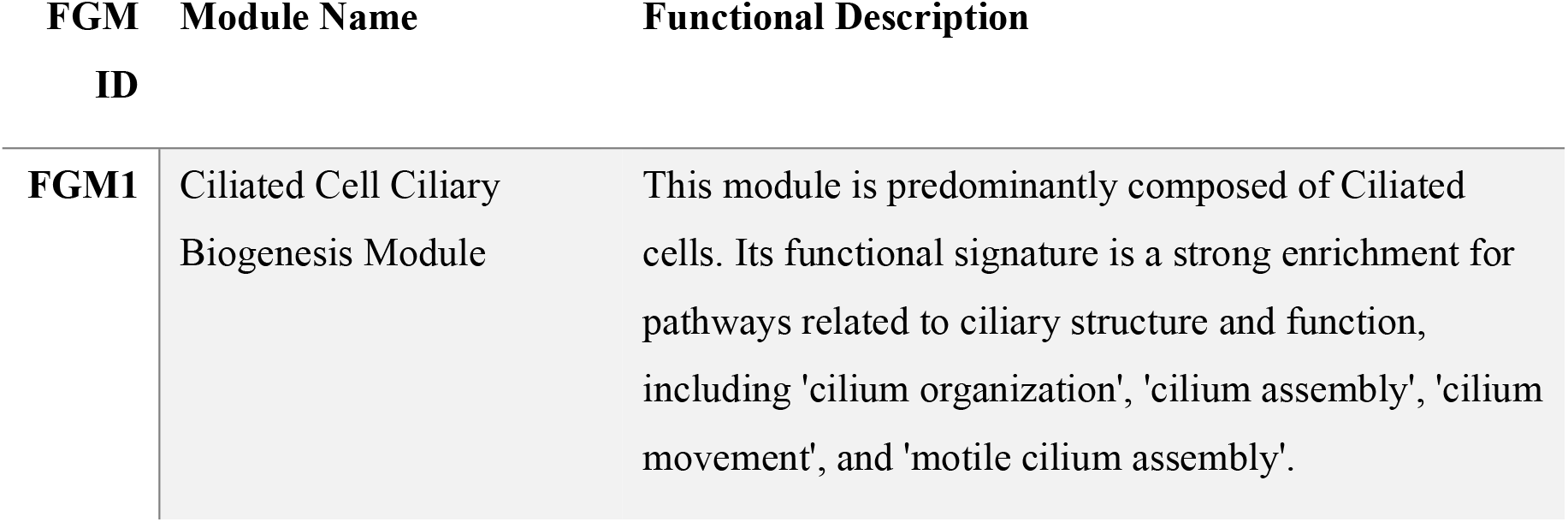
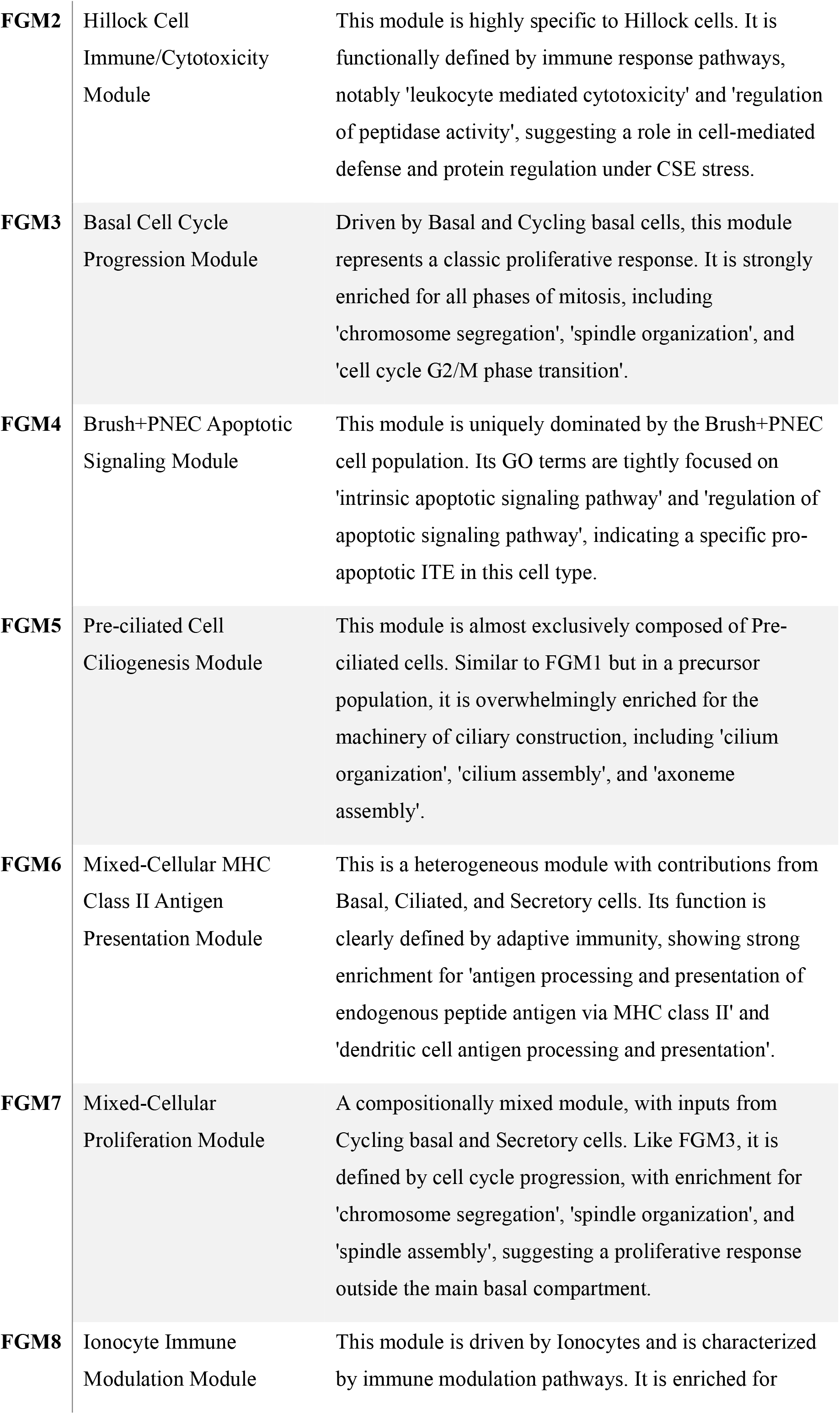
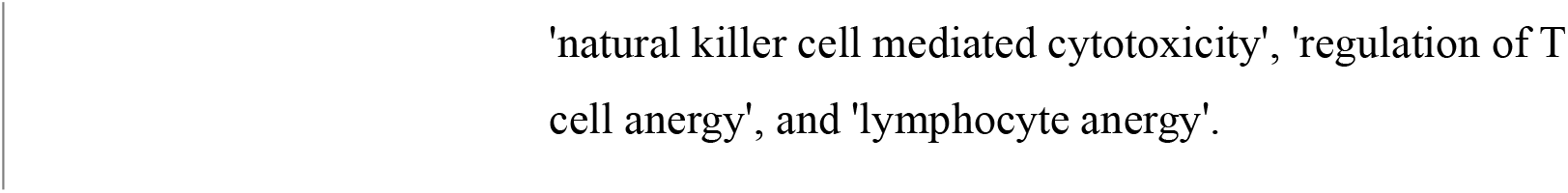
Details on the FGM identified in primary human bronchial organoids under cigarette-smoke extract (CSE) exposure.

**Supplementary Table S4.** Details on the FGM identified in Kang dataset.

| **FGM NO.** | **SHORT NAME** | **DETAILED FUNCTIONAL DESCRIPTION (GO ENRICHMENT SUMMARY)** |
| --- | --- | --- |
| **FGM 1** | MHC I/II Presentation & T Cell Regulation | Highly enriched in MHC I/II antigen processing and presentation (both classes) and the Regulation of leukocyte and T cell proliferation. Also includes positive regulation of Type II IFN production. |
| **FGM 2** | Apoptosis & Metabolic Negative Regulation | Centered on the Regulation of apoptotic signaling pathway and Negative regulation of metabolic processes, including purine nucleotide catabolism and glycolysis, often via mitochondrial pathways. |
| **FGM 3** | Antiviral Defense & Lymphocyte Differentiation | Core function is the Defense response to virus and Lymphocyte differentiation. Key features include Viral genome replication regulation and Granzyme-mediated programmed cell death signaling. |
| **FGM 4** | Protein Folding & Heat Stress Response | Highly specialized for Protein folding ('de novo', chaperone-mediated) and the Response to heat or temperature stimulus. Reflects the cellular machinery maintaining proteostasis. |
| **FGM 5** | T Cell Activation & Antiviral Immunity | Characterized by a strong Defense response to virus and the Alpha-beta T cell activation and differentiation, including negative regulatory aspects of these processes. |
| **FGM 6** | MHC II Antigen Presentation (DC/Core) | Highly specific for the Antigen processing and presentation of exogenous peptide via MHC class II. Also includes MHC class II protein complex assembly and Positive regulation of T cell activation. |
| **FGM 7** | MHC II Antigen Presentation | Functional profile is nearly identical to FGM6, focusing on Antigen processing and presentation of exogenous antigen via |
|  | (B Cell/Core) | MHC class II, confirming this as a central APC response. |
| **FGM 8** | Immune Cell Migration & Radiation Stress | Defined by Mononuclear/Lymphocyte/T cell migration and chemotaxis. Also includes the Response to gamma and ionizing radiation and Positive regulation of calcium ion release. |

**Supplementary Table S5.**
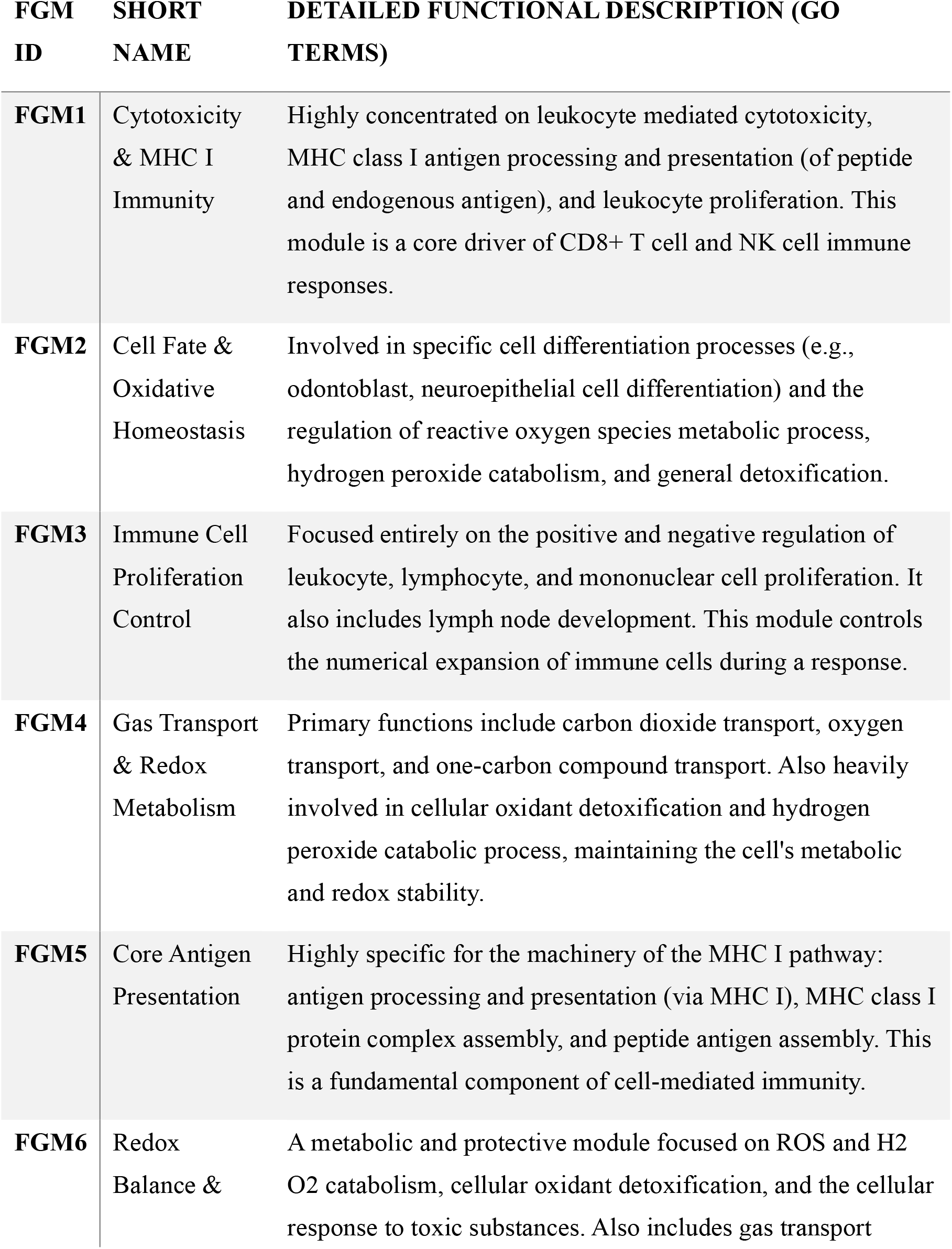
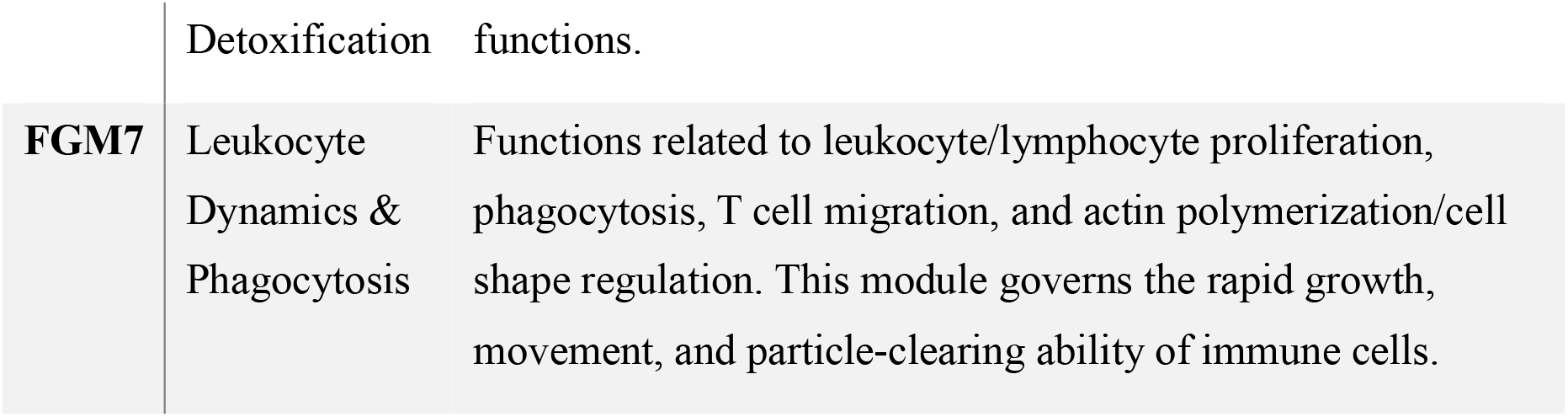
Details on the FGM identified in Norman dataset.

## References

[1] Dixit, A., Parnas, O., Li, B., Chen, J., Fulco, C.P., Jerby-Arnon, L., Marjanovic, N.D., Dionne, D., Burks, T., Raychowdhury, R., et al.: Perturb-seq: dissecting molecular circuits with scalable single-cell rna profiling of pooled genetic screens. Cell 167(7), 1853–1866 (2016)

[2] Srivatsan, S.R., McFaline-Figueroa, J.L., Ramani, V., Saunders, L., Cao, J., Packer, J., Pliner, H.A., Jackson, D.L., Daza, R.M., Christiansen, L., et al.: Massively multiplex chemical transcriptomics at single-cell resolution. Science 367(6473), 45–51 (2020)

[3] Peidli, S., Green, T.D., Shen, C., Gross, T., Min, J., Garda, S., Yuan, B., Schumacher, L.J., Taylor-King, J.P., Marks, D.S., et al.: scPerturb: harmonized single-cell perturbation data. Nature Methods 21(3), 531–540 (2024)

[4] Lotfollahi, M., Wolf, F.A., Theis, F.J.: scGen predicts single-cell perturbation responses. Nature Methods 16(8), 715–721 (2019)

[5] Adduri, A.K., Gautam, D., Bevilacqua, B., Imran, A., Shah, R., Naghipourfar, M., Teyssier, N., Ilango, R., Nagaraj, S., Dong, M., et al.: Predicting cellular responses to perturbation across diverse contexts with state. bioRxiv, 2025–06 (2025)

[6] Dong, M., Wang, B., Wei, J. O. Fonseca, A.H., Perry, C.J., Frey, A., Ouerghi, F., Foxman, E.F., Ishizuka, J.J., Dhodapkar, R.M., et al.: Causal identification of single-cell experimental perturbation effects with cinema-ot. Nature Methods 20(11), 1769–1779 (2023)

[7] Chewi, S., Niles-Weed, J., Rigollet, P.: Statistical optimal transport (2024)

[8] Fukumizu, K., Bach, F.R., Jordan, M.I.: Dimensionality reduction for supervised learning with reproducing kernel hilbert spaces. Journal of Machine Learning Research 5(Jan), 73–99 (2004)

[9] Sheng, T., Sriperumbudur, B.K.: On distance and kernel measures of conditional dependence. Journal of Machine Learning Research 24(7), 1–16 (2023)

[10] Louizos, C., Welling, M., Kingma, D.P.: Learning sparse neural networks through l 0 regularization. In: International Conference on Learning Representations (2018)

[11] Khemakhem, I., Kingma, D., Monti, R., Hyvarinen, A.: Variational autoen-coders and nonlinear ica: A unifying framework. In: International Conference on Artificial Intelligence and Statistics, pp. 2207–2217 (2020). PMLR

[12] Zheng, Y., Zhang, K.: Generalizing nonlinear ica beyond structural sparsity. Advances in Neural Information Processing Systems 36, 13326–13355 (2023)

[13] Balakrishnan, S., Kennedy, E., Wasserman, L.: Conservative inference for counterfactuals. Journal of Causal Inference 13(1), 20230071 (2025)

[14] Kang, H.M., Subramaniam, M., Targ, S., Nguyen, M., Maliskova, L., McCarthy, E., Wan, E., Wong, S., Byrnes, L., Lanata, C.M., et al.: Multiplexed droplet single-cell rna-sequencing using natural genetic variation. Nature Biotechnology 36(1), 89–94 (2018)

[15] Hagai, T., Chen, X., Miragaia, R.J., Rostom, R., Gomes, T., Kunowska, N., Henriksson, J., Park, J.-E., Proserpio, V., Donati, G., et al.: Gene expression variability across cells and species shapes innate immunity. Nature 563(7730), 197–202 (2018)

[16] Norman, T.M., Horlbeck, M.A., Replogle, J.M., Ge, A.Y., Xu, A., Jost, M., Gilbert, L.A., Weissman, J.S.: Exploring genetic interaction manifolds constructed from rich single-cell phenotypes. Science 365(6455), 786–793 (2019)

[17] Gagliardi, T.B., Goldstein, M.E., Song, D., Gray, K.M., Jung, J.W., Ignacio, M.A., Stroka, K.M., Duncan, G.A., Scull, M.A.: Rhinovirus c replication is associated with the endoplasmic reticulum and triggers cytopathic effects in an in vitro model of human airway epithelium. PLoS pathogens 18(1), 1010159 (2022)

[18] Brekman, A., Walters, M.S., Tilley, A.E., Crystal, R.G.: Foxj1 prevents cilia growth inhibition by cigarette smoke in human airway epithelium in vitro. American journal of respiratory cell and molecular biology 51(5), 688–700 (2014)

[19] Crystal, R.G.: Airway basal cells. the “smoking gun” of chronic obstructive pulmonary disease. American journal of respiratory and critical care medicine 190(12), 1355–1362 (2014)

[20] Lan, M.-Y., Ho, C.-Y., Lee, T.-C., Yang, A.-H.: Cigarette smoke extract induces cytotoxicity on human nasal epithelial cells. American journal of rhinology 21(2), 218–223 (2007)

[21] Gong, Y., Xu, J., Wu, M., Gao, R., Sun, J., Yu, Z., Zhang, Y.: Single-cell biclustering for cell-specific transcriptomic perturbation detection in ad progression. Cell Reports Methods 4(4) (2024)

[22] Xu, S., Hu, E., Cai, Y., Xie, Z., Luo, X., Zhan, L., Tang, W., Wang, Q., Liu, B., Wang, R., et al.: Using clusterprofiler to characterize multiomics data. Nature Protocols 19(11), 3292–3320 (2024)

[23] Crouse, J., Bedenikovic, G., Wiesel, M., Ibberson, M., Xenarios, I., Von Laer, D., Kalinke, U., Vivier, E., Jonjic, S., Oxenius, A.: Type i interferons protect t cells against nk cell attack mediated by the activating receptor ncr1. Immunity 40(6), 961–973 (2014)

[24] Pramanik, J., Chen, X., Kar, G., Henriksson, J., Gomes, T., Park, J.-E., Natarajan, K., Meyer, K.B., Miao, Z., McKenzie, A.N., et al.: Genome-wide analyses reveal the ire1a-xbp1 pathway promotes t helper cell differentiation by resolving secretory stress and accelerating proliferation. Genome Medicine 10(1), 76 (2018)

[25] Dahlgren, M.W., Plumb, A.W., Niss, K., Lahl, K., Brunak, S., Johansson-Lindbom, B.: Type i interferons promote germinal centers through b cell intrinsic signaling and dendritic cell dependent th1 and tfh cell lineages. Frontiers in Immunology 13, 932388 (2022)

[26] Li, Q., Wu, Y., Zhang, Y., Sun, H., Lu, Z., Du, K., Fang, S., Li, W.: mir-125b regulates cell progression in chronic myeloid leukemia via targeting bak1. American Journal of Translational Research 8(2), 447 (2016)

[27] Tallack, M.R., Perkins, A.C.: Klf1 directly coordinates almost all aspects of terminal erythroid differentiation. IUBMB Life 62(12), 886–890 (2010)

[28] Ianniciello, A., Zarou, M.M., Rattigan, K.M., Scott, M., Dawson, A., Dunn, K., Brabcova, Z., Kalkman, E.R., Nixon, C., Michie, A.M., et al.: Ulk1 inhibition promotes oxidative stress–induced differentiation and sensitizes leukemic stem cells to targeted therapy. Science Translational Medicine 13(613), 5016 (2021)

[29] Shyamsunder, P., Shanmugasundaram, M., Mayakonda, A., Dakle, P., Teoh, W.W., Han, L., Kanojia, D., Lim, M.C., Fullwood, M., An, O., et al.: Identification of a novel enhancer of cebpe essential for granulocytic differentiation. Blood, The Journal of the American Society of Hematology 133(23), 2507–2517 (2019)

[30] Kaweme, N.M., Zhou, S., Changwe, G.J., Zhou, F.: The significant role of redox system in myeloid leukemia: from pathogenesis to therapeutic applications. Biomarker Research 8(1), 63 (2020)

[31] Ramirez-Carrozzi, V.R., Braas, D., Bhatt, D.M., Cheng, C.S., Hong, C., Doty, K.R., Black, J.C., Hoffmann, A., Carey, M., Smale, S.T.: A unifying model for the selective regulation of inducible transcription by cpg islands and nucleosome remodeling. Cell 138(1), 114–128 (2009)

[32] Fenouil, R., Cauchy, P., Koch, F., Descostes, N., Cabeza, J.Z., Innocenti, C., Ferrier, P., Spicuglia, S., Gut, M., Gut, I., et al.: Cpg islands and gc content dictate nucleosome depletion in a transcription-independent manner at mammalian promoters. Genome research 22(12), 2399–2408 (2012)

[33] Masoudi-Sobhanzadeh, Y., Li, S., Peng, Y., Panchenko, A.R.: Interpretable deep residual network uncovers nucleosome positioning and associated features. Nucleic Acids Research 52(15), 8734–8745 (2024)

[34] Schwab, A., Siddiqui, A., Vazakidou, M.E., Napoli, F., Böttcher, M., Menchicchi, B., Raza, U., Saatci, Ö., Krebs, A.M., Ferrazzi, F., et al.: Polyol pathway links glucose metabolism to the aggressiveness of cancer cells. Cancer Research 78(7), 1604–1618 (2018)

[35] Chen, X., Chen, C., Hao, J., Qin, R., Qian, B., Yang, K., Zhang, J., Zhang, F.: Akr1b1 upregulation contributes to neuroinflammation and astrocytes proliferation by regulating the energy metabolism in rat spinal cord injury. Neurochemical Research 43(8), 1491–1499 (2018)

[36] Batchuluun, B., Pinkosky, S.L., Steinberg, G.R.: Lipogenesis inhibitors: therapeutic opportunities and challenges. Nature Reviews Drug Discovery 21(4), 283–305 (2022)

[37] Sun, J., Liang, C., Wei, R., Zheng, P., Bai, L., Ouyang, W., Yan, H., Ye, P.: scmrdr: A scalable and flexible framework for unpaired single-cell multi-omics data integration. arXiv preprint arXiv:2510.24987 (2025)

[38] Tejada-Lapuerta, A., Bertin, P., Bauer, S., Aliee, H., Bengio, Y., Theis, F.J.: Causal machine learning for single-cell genomics. Nature Genetics, 1–12 (2025)

[39] Klein, D., Palla, G., Lange, M., Klein, M., Piran, Z., Gander, M., Meng-Papaxanthos, L., Sterr, M., Saber, L., Jing, C., et al.: Mapping cells through time and space with moscot. Nature 638(8052), 1065–1075 (2025)

[40] Metzner, E., Southard, K.M., Norman, T.M.: Multiome perturb-seq unlocks scalable discovery of integrated perturbation effects on the transcriptome and epigenome. Cell systems 16(1) (2025)

[41] Lopez, R., Regier, J., Cole, M.B., Jordan, M.I., Yosef, N.: Deep generative modeling for single-cell transcriptomics. Nature Methods 15(12), 1053–1058 (2018)

[42] Higgins, I., Matthey, L., Pal, A., Burgess, C., Glorot, X., Botvinick, M., Mohamed, S., Lerchner, A.: beta-VAE: Learning basic visual concepts with a constrained variational framework. In: International Conference on Learning Representations (2017)

[43] Lopez, R., Regier, J., Jordan, M.I., Yosef, N.: Information constraints on auto-encoding variational bayes. Advances in Neural Information Processing Systems 31 (2018)

[44] Fukumizu, K., Gretton, A., Sun, X., Schölkopf, B.: Kernel measures of conditional dependence. Advances in neural information processing systems 20 (2007)

[45] Zheng, Y., Ng, I., Zhang, K.: On the identifiability of nonlinear ica: Sparsity and beyond. Advances in Neural Information Processing Systems 35, 16411–16422 (2022)

[46] Turner, H., Bailey, T., Krzanowski, W.: Improved biclustering of microarray data demonstrated through systematic performance tests. Computational Statistics & Data Analysis 48(2), 235–254 (2005)

[47] Gretton, A., Borgwardt, K.M., Rasch, M.J., Schölkopf, B., Smola, A.: A kernel two-sample test. The Journal of Machine Learning Research 13(1), 723–773 (2012)

## References

[9] Ramirez-Carrozzi, V. R., Braas, D., Bhatt, D. M., Cheng, C. S., Hong, C., Doty, K. R., … & Smale, S. T. (2009). A unifying model for the selective regulation of inducible transcription by CpG islands and nucleosome remodeling. Cell, 138(1), 114–128.

[10] Ahmad, K., Henikoff, S., & Ramachandran, S. (2022). Managing the steady state chromatin landscape by nucleosome dynamics. Annual review of biochemistry, 91(1), 183–195.

[11] Fenouil, R., Cauchy, P., Koch, F., Descostes, N., Cabeza, J. Z., Innocenti, C., … & Andrau, J. C. (2012). CpG islands and GC content dictate nucleosome depletion in a transcription-independent manner at mammalian promoters. Genome research, 22(12), 2399–2408.

[12] Masoudi-Sobhanzadeh, Y., Li, S., Peng, Y., & Panchenko, A. R. (2024). Interpretable deep residual network uncovers nucleosome positioning and associated features. Nucleic Acids Research, 52(15), 8734–8745.

## Reference

[1] Zheng, Y., Ng, I., & Zhang, K. (2022). On the identifiability of nonlinear ICA: Sparsity and beyond. Advances in neural information processing systems, 35, 16411–16422.

[2] Zheng, Y., & Zhang, K. (2023). Generalizing nonlinear ICA beyond structural sparsity. Advances in Neural Information Processing Systems, 36, 13326–13355.

[3] Li, Z., Fan, S., Zheng, Y., Ng, I., Xie, S., Chen, G., … & Zhang, K. (2025). Synergy Between Sufficient Changes and Sparse Mixing Procedure for Disentangled Representation Learning. In The Thirteenth International Conference on Learning Representations.

[4] Khemakhem, I., Kingma, D., Monti, R., & Hyvarinen, A. (2020). Variational autoencoders and nonlinear ica: A unifying framework. In International conference on artificial intelligence and statistics (pp. 2207–2217). PMLR.

[5] Louizos, C., Welling, M., & Kingma, D. P. (2018). Learning Sparse Neural Networks through L0 Regularization. In International Conference on Learning Representations.

[6] Xie, S., Kong, L., Gong, M., & Zhang, K. (2023). Multi-domain image generation and translation with identifiability guarantees. In The Eleventh international conference on learning representations.

[7] Chewi, S., Niles-Weed, J., & Rigollet, P. (2024). Statistical optimal transport. Springer.

[8] Balakrishnan, S., Kennedy, E., & Wasserman, L. (2025). Conservative inference for counterfactuals. Journal of Causal Inference, 13(1), 20230071.

